# Fibronectin receptor integrin α5β1 regulates assembly of PP2A complexes through PDE4D: modulation of vascular inflammation and atherosclerosis

**DOI:** 10.1101/680728

**Authors:** Sanguk Yun, Rui Hu, Melanie E. Schwaemmle, Alexander N. Scherer, Zhenwu Zhuang, Anthony J. Koleske, David C. Pallas, Martin A. Schwartz

**Affiliations:** Department of Internal Medicine, Yale Cardiovascular Research Center, Yale University, New Haven, CT 06520, USA; Department of Molecular Biophysics and Biochemistry, Yale University, New Haven, CT 06520, USA; Department of Biochemistry, Emory University School of Medicine, 1510 Clifton Rd., Atlanta, GA 30322, USA; Department of Biomedical Engineering, Yale University, New Haven, CT 06520, USA; Department of Cell Biology, Yale University, New Haven, CT 06520, USA

## Abstract

Fibronectin in the vascular wall promotes inflammatory activation of the endothelium during vascular remodeling and atherosclerosis. These effects are mediated in part by fibronectin binding to integrin α5, which recruits and activates phosphodiesterase 4D5 (PDE4D5) by inducing its dephosphorylation on an inhibitory site Ser651. Active PDE then hydrolyzes anti-inflammatory cAMP to facilitate inflammatory signaling. To test this model in vivo, we mutated the integrin binding site in PDE4D5 in mice. This mutation reduced endothelial inflammatory activation in athero-prone regions of arteries, and, in a hyperlipidemia model, reduced atherosclerotic plaque size while increasing markers of plaque stability. We then investigated the mechanism of PDE4D5 activation. Proteomics identified the PP2A regulatory subunit B55α as the factor recruiting PP2A to PDE4D5. The B55α-PP2A complex localized to adhesions and directly dephosphorylated PDE4D5. This interaction also unexpectedly stabilized the PP2A-B55α complex. The integrin-regulated, pro-atherosclerotic transcription factor Yap is also dephosphorylated and activated through this pathway. PDE4D5 therefore mediates matrix-specific regulation of EC phenotype via an unconventional adapter role, assembling and anchoring a multifunctional PP2A complex with other targets. These results are likely to have widespread consequences for control of cell function by integrins.

## Introduction

Endothelial basement membranes in stable vessels, consisting primarily of collagen IV, laminins and less abundant species, promote vessel maturation and stability (1, 2). By contrast, fibronectin (FN) is present at low levels in stable vessels in most tissues but is strongly upregulated in inflammation, developmental and postnatal angiogenesis, and in atherosclerosis, as is its main receptor integrin α5β1 (3–6). FN has been most studied in atherogenesis, where it contributes to plaque formation (7). These effects are mediated in part by inhibiting the anti-inflammatory cAMP/protein kinase A (PKA) pathway (8). This occurs through binding of the α5 subunit cytoplasmic domain to the cAMP-specific phosphodiesterase PDE4D5, which results in dephosphorylation of PDE4D5 on an inhibitory site (9). The resultant increase in PDE catalytic activity suppresses cAMP/PKA signaling, thus priming ECs for inflammatory activation.

EC responses to fluid shear stress play major roles in vessel function, remodeling and disease. Shear stress regulates artery remodeling to determine lumen diameter and, during angiogenesis, induces stabilization of vessel sprouts once flow is re-established (10, 11). Disturbed flow patterns that arise in regions of arteries that curve sharply or branch induce local EC inflammatory activation. In the presence of systemic risk factors such as hyperlipidemia, hyperglycemia, hypertension and elevated inflammatory mediators, these regions selectively develop atherosclerotic plaques. FN gene expression and matrix assembly are induced by disturbed flow (12, 13). FN is deposited at athero-prone regions of arteries in wild type (WT) mice and increases in atherosclerotic lesions in mice and humans (4, 14). Deletion of each of the individual isoforms of FN reduces plaque size in mice (15–17). However, in the one study where it was examined, deletion of plasma FN also resulted in thinner fibrous caps with evidence of plaque rupture, suggesting reduced plaque stability (17).

Adherence of cells to FN promoted activation of NF-κB and other inflammatory pathways in response to disturbed flow, IL-1β and oxidized LDL, compared to adherence to collagens or laminins (4, 9, 18, 19). These effects were reversed by mutation of the integrin α5 cytoplasmic domain in vitro. To test in vivo, we introduced this mutation into mice, where it suppressed atherosclerosis and improved recovery from hindlimb ischemia (9) (20). EC-specific deletion of integrin α5 also strongly reduced atherosclerosis in mouse models (21). However, other integrin α5 cytoplasmic domain effectors may also mediate these effects (22).

We set out to rigorously test to what extent binding of PDE4D5 to integrin α5 mediates α5’s role in vascular inflammation and atherosclerosis. Because complete deletion of PDE4D results in neonatal growth retardation and lethality (23), we mutated the integrin binding sequence in PDE4D5 to block this interaction without interfering with other functions. Positive results then prompted us to further examine the mechanism of PDE4D5 regulation. These studies revealed that PDE4D5 regulates a wider array of pathways via effects on phosphatase PP2A and other downstream targets.

## Results

### PDE4D^mut^ mice

We previously mapped the integrin α5 binding site in PDE4D5 to a short basic sequence (KKKR) in the linker region between the N-terminal UCR2 domain and the catalytic domain (9). Replacement of this sequence with ‘EEEE’ abolished binding to integrin α5 and blocked the pro-inflammatory effect of FN in vitro. In this study, we aimed to probe its function in vivo by engineering this mutation into mice using CRISPR/Cas9 technology (Fig 1A). To minimize off-target mutations, we used the double nicking strategy with Cas9 nickase and two guide RNAs that recognize flanking target sequences (24) (Fig. 1A). From 15 pups, sequencing and genotyping identified one mouse with the correct mutation (Fig. 1B, Fig. S1A; 1 female founder, 3 in-del mutants and 11 wild type mice). The mutated allele was successfully transmitted to the next generation (Fig. S1B). Homozygous mice were born at the expected Mendelian ratio (WT: 28, Het: 45, Homo: 20) and were viable and fertile, without obvious abnormalities. PDE4D expression was similar in the PDE4D^mut^ compared to wild type mice (Fig. S1C).

**Figure 1.**
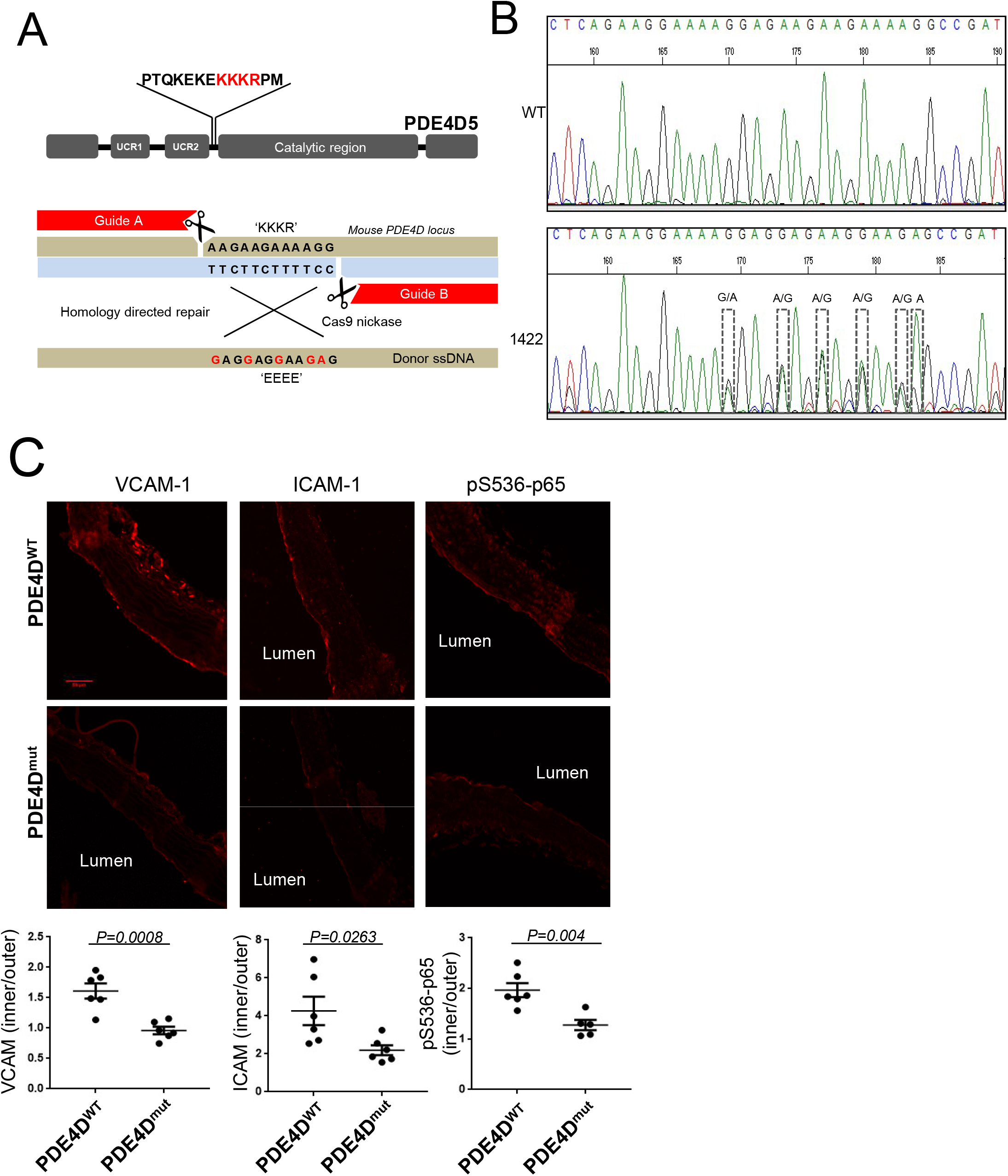
PDE4D^mut^ mice showed reduced inflammation in a region of disturbed flow. **A.** Gene editing strategy. Two guide RNA’s with Cas9 nickase were used with donor ssDNA to induce the KKKR to EEEE mutation. **B.** Confirming the mutation. The female founder mouse (1422) identified by PCR with mutant-specific primers in Fig S1A was confirmed by sequencing of genomic DNA. **C.** Inflammatory markers. Longitudinal sections from aortas from three month old wild type or mutant mice were stained for the indicated inflammatory markers. Images show the inner curvature of the aortic arch. Intensity for each marker in the endothelial layer was quantified as described in Methods and intensity in the inner curvature normalized by signal from the outer curvature. Values are means ± SEM. P value determined by two-tailed t-tests.

ECs in atherosclerosis-prone regions of arteries in WT mice show elevated inflammatory markers that coincide with FN accumulation (4, 9). To investigate effects of the PDE4D5 mutation, longitudinal section of aortas from three-month-old mice were examined for inflammatory markers in the endothelium in the inner curvature of the aortic arch, a well characterized region of disturbed flow that is prone to atherosclerosis when hyperlipidemia and other risk factors are present. As expected, ICAM-1, VCAM-1 and phosphorylated NFκB p65 were all elevated in this region in WT mice (quantified as the ratio of staining relative to the athero-resistant outer curvature in each mouse); these markers were all significantly (ICAM-1) or completely (NF-κB and VCAM-1) suppressed in PDE4D^mut^ mice (Fig. 1C). Preventing the binding of PDE4D5 to integrin α5 thus inhibits vascular inflammation similarly to mutation of the α5 cytoplasmic domain (25).

Next, PDE4D^mut^ mice were bred onto the ApoE null background and fed a high fat, “Western” diet for four months to induce atherosclerotic lesions. The PDE mutation had no effect on serum lipids (Fig. S2A). Oil red-O staining of aortic roots showed that atherosclerotic lesions were significantly reduced in PDE4D^mut^/ApoE null compared to WT/ApoE null mice (Fig. 2A). CD68 staining for macrophages was similarly reduced (Fig. 2B). Previous studies showed evidence of plaque vulnerability after deletion of plasma FN (26). However, smooth muscle actin, quantified per plaque area, was increased by 50% in PDE4D^mut^ mice (Fig. 2F); picrosirius red staining for fibrillar collagen was also higher in PDE4D^mut^ mice (Fig. 2C). By contrast, MMP-2 and MMP-9 per area of plaque were reduced (Fig 2D, E), as was MMP activity (Fig. S2C). Together, these data show that mutation of the integrin binding site in PDE4D5 reduces atherosclerotic lesion size and inflammatory status in hyperlipidemic mice while increasing markers of plaque stability. These results provide definitive evidence that PDE4D5 is a critical integrin α5 effector in this pathway.

**Figure 2.**
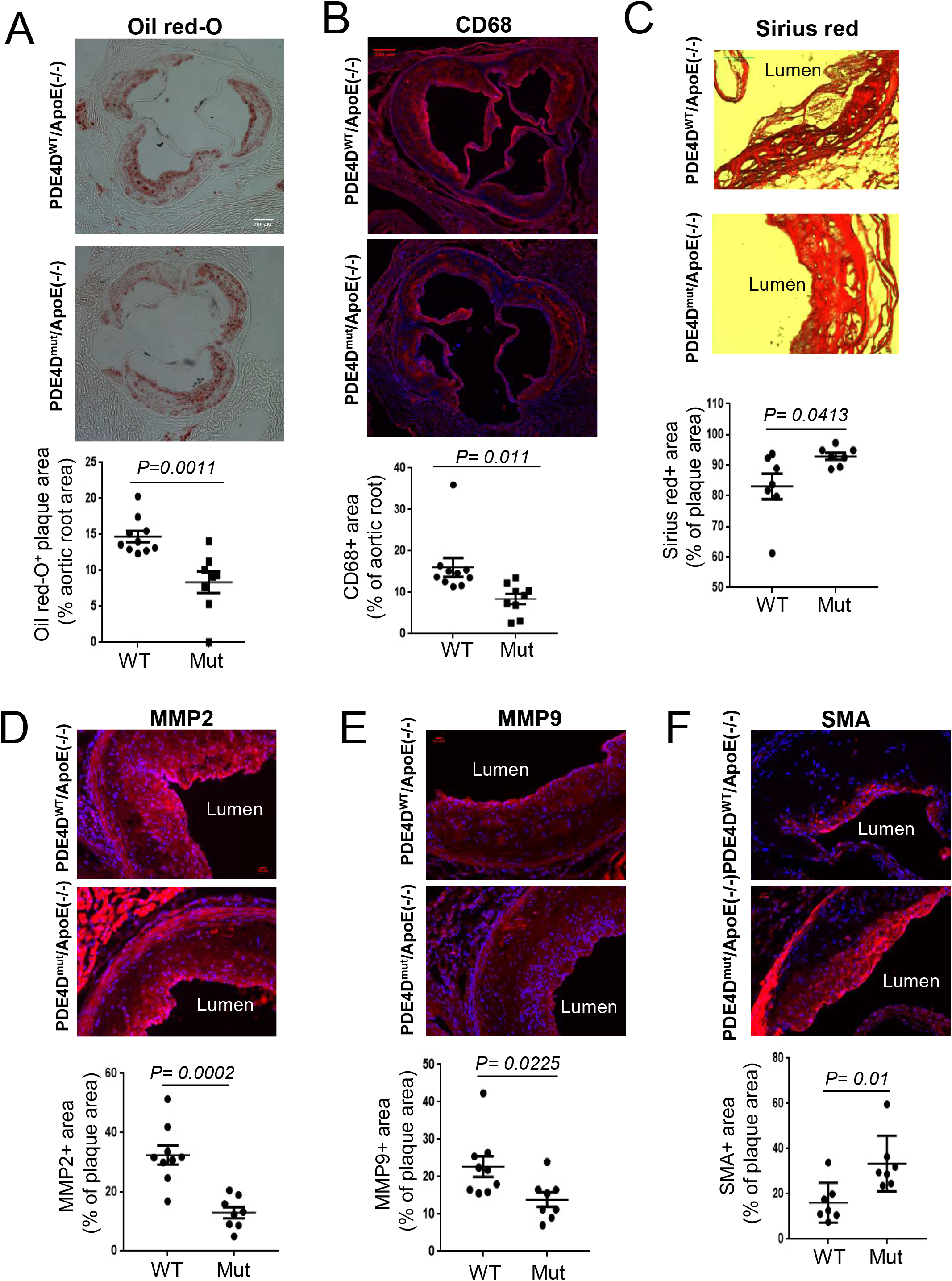
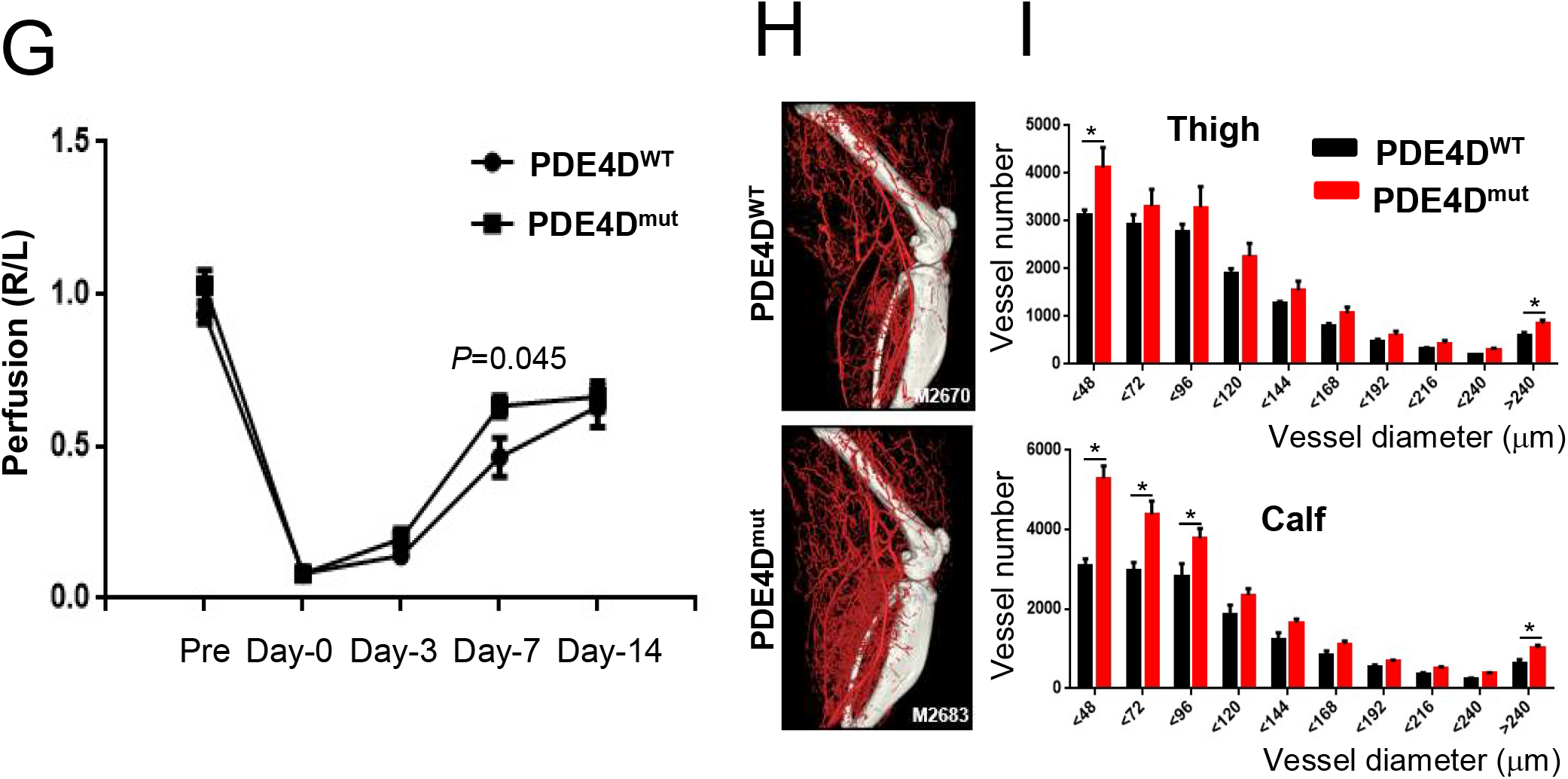
Atherosclerosis and recovery from hindlimb ischemia. **A.** Lesion size. Aortic root sections from WT or PDE4D^mut^ ApoE null mice after 4 months on a high fat diet were stained with Oil Red-O. **B.** Macrophage content. Section were stained for CD68 to identify macrophages. **C-E.** Plaque composition. Sections were stained for fibrillar collagens with Picrosirius Red (C), MMP 2 (D) and MMP 9 (E). **F.** Smooth muscle actin. Sections were stained for SMA to identify smooth muscle cells. In all cases, staining intensity was quantified as described in Methods and normalized to total plaque area. **G.** Blood flow recovery. Femoral artery ligation was performed on the right leg of three month old wild type or mutant mice and blood flow measured by laser Doppler at each time; values are means ±SEM, n=6. **H, I.** MicroCT of ischemic legs 14 days after surgery. Results were quantified according to vessel diameter. * p< 0.05 in two-tailed t-tests.

Recovery from restriction of arterial flow occurs via both flow-dependent arteriogenesis and ischemia/VEGF-induced angiogenesis (27, 28). As FN has been implicated in both processes (5, 6, 28, 29), we tested PDE4D^mut^ mice in the hindlimb ischemia model where ligation of the femoral artery induces ischemia-independent flow-dependent arteriogenesis in the upper leg and ischemia-induced angiogenesis in the lower leg (30). Three month old wild type or PDE4D^mut^ mice were subjected to femoral artery ligation and blood flow recovery was monitored by laser Doppler imaging. Blood flow recovery was modestly but significantly enhanced in PDE4D^mut^ mice 7 days after ligation (Figure 2G, Figure S2D). MicroCT showed that PDE4D^mut^ mice had higher density of both large and small vessels in the ischemic legs 14 days after surgery (Figure 2H, I). No differences were observed in vessel density in the control legs (Figure S2E, F), indicating that these effects occurred only after surgery. Specifically blocking the PDE4D5-integrin interaction therefore did not limit but instead slightly accelerated vessel remodeling in C57Bl6 mice.

### Identification of the PP2A complex that dephosphorylates PDE4D

As PDE4D5 activation appeared to be critical for these effects, we next investigated the mechanism of its activation in more detail. Integrin α5β1 activates PDE4D5 by triggering its dephosphorylation on inhibitory S^651^ by PP2A, which suppresses anti-inflammatory cAMP-PKA signaling (9). PP2A however represents a family of enzyme complexes comprised of a catalytic subunit (C), scaffolding subunit (A) and regulatory/targeting subunit (B). There are two C isoforms, two A isoforms, and at least 23 regulatory B subunits that mediate selective substrate binding, subcellular localization and catalytic activity of PP2A holoenzymes (31). PP2A AC-core dimers dynamically associate with B subunits regulated by posttranslational modification such as phosphorylation and methylation of C subunit or by adaptor protein binding (32, 33).

To identify the specific PP2A complex mediating FN-dependent PDE4D dephosphorylation, we hypothesized that phospho-mimetic mutant (PDE4D5-S651E) would behave as a phosphatase trapping mutant for affinity chromatography. Cells were therefore transfected with GFP-PDE4D5-S651E, detached from the culture dishes and replated on dishes coated with FN or with laminin/coll IV basement membrane protein (matrigel, MG). Lysates were pulled down with GFP-Trap nanobody beads and analyzed by SDS-PAGE and silver staining. We observed a PDE4D5-dependent band at 55kD from cells on FN but not MG (Fig. 3A). LC-MS proteomics identified the protein as the PP2A-B55α subunit (Fig S3). This finding was confirmed by immunoblotting (Fig. 3B). Thus, B55α appears to be a FN-specific PP2A regulatory/targeting subunit that binds PDE4D5.

**Figure 3.**
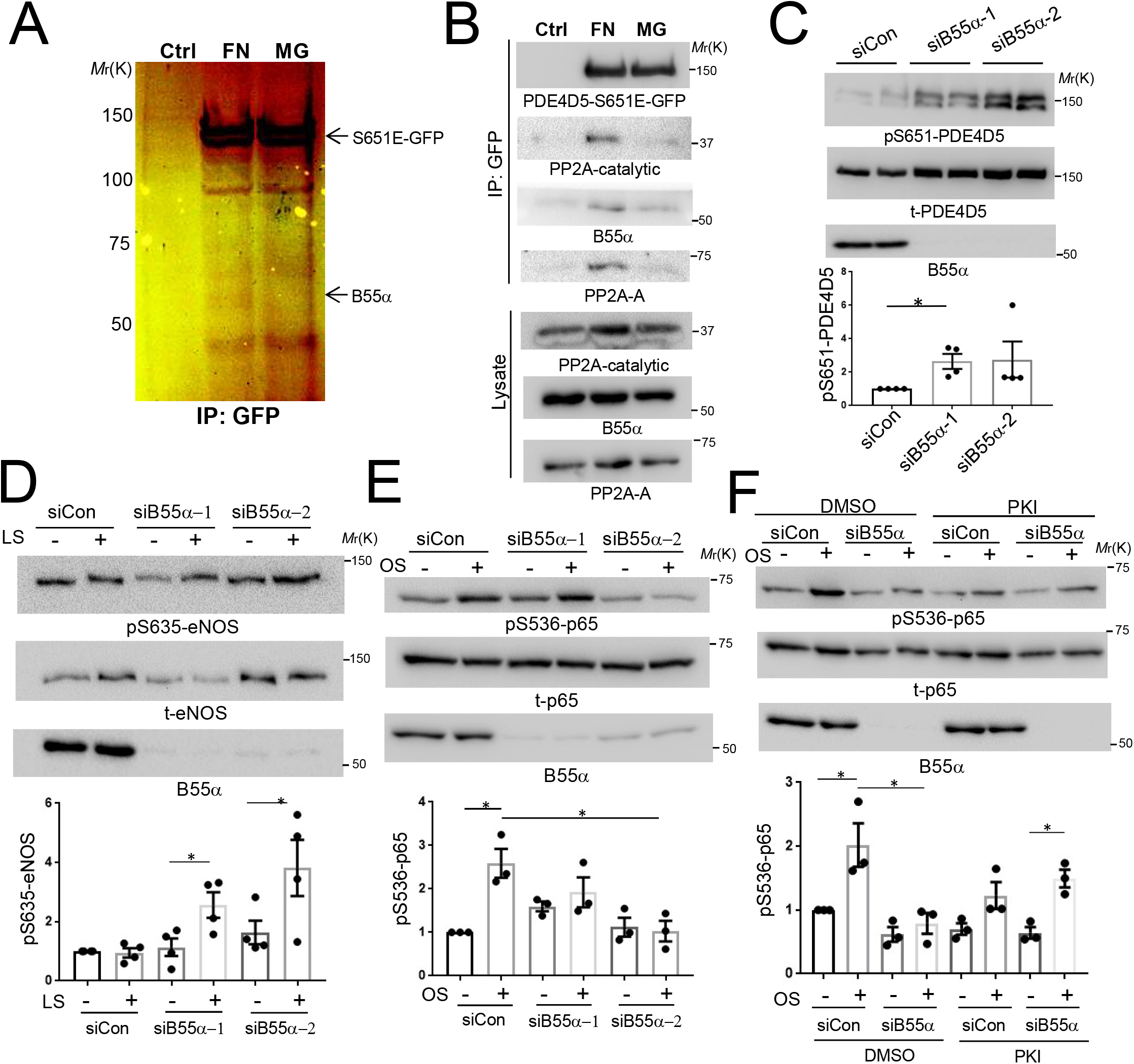
B55a is required for PP2A-dependent PDE4D dephosphorylation. **A.** FN promotes B55α binding to PDE4D5. BAECs stably expressing GFP-tagged PDE4D5-S651E were plated on dishes coated with FN or matrigel (MG) for 20 min. Lysates were immunoprecipitated with GFP-Trap. SDS PAGE and silver staining identified a.FN-specific band at ~55kDa, which was identified by LC-MS (Fig. S3) as B55α. **B.** Confirming proteomic results. PDE4D5-S651E immune complexes prepared as in A were Western blotted for PP2A subunits as indicated. Results are representative of three independent experiments. **C.** B55α knockdown. BAECs expressing WT GFP-tagged PDE4D5 were transfected with control or B55α siRNAs and plated on FN for 6 hr. Cell lysates were Western blotted for PDE4D S^651^ phosphorylation or total PDE4D (n=4). **D.** BAECs transfected with control or B55α siRNAs were plated on FN and exposed to laminar shear (LS) for 15 min. Flow-dependent S^635^ phosphorylation of eNOS was measured by immunoblotting with phospho-specific antibody (n=4). **E.** BAECs transfected with control or B55α siRNAs were plated on FN and subjected to oscillatory shear (OS) for 2 hr. NF-kB activation was measured by Western blotting for p65 S^536^ phosphorylation (n=3). **F.** BAECs transfected with control or B55α siRNA were plated on FN, treated with or without the PKI 14-22 amide inhibitor (PKI) or dimethylsulfoxide (DMSO), and subjected to OS for 2 hr. NF-kB activation was measured by probing for p65 pS^536^ (n=3). *p<0.05 by two tailed t-test (C, D) or one way ANOVA (E, F)

To test its function in this pathway, we knocked down B55α in ECs using two different siRNAs. B55α depletion strongly increased PDE4D5-S^651^ phosphorylation, supporting its role in PDE4D5 dephosphorylation (Fig. 3C). To test the role of cAMP and PKA, we examined the specific PKA site (S^635^) on eNOS (34). B55α was therefore knocked down in ECs, which were plated on FN and stimulated by flow. eNOS S635 phosphorylation is normally suppressed by FN (19) but was increased following B55α depletion (Fig 3D). Consistent with high cAMP/PKA signaling, NF-κB activation by oscillatory shear stress (OS) was also inhibited (Fig. 3E). To test whether the reduced NF-κB activation was mediated by PKA, cells were treated with the specific PKA inhibitor, myristoylated PKI (14–22) amide. This treatment restored flow-dependent NFκB activation after B55α depletion (Fig. 3F). Together, these results show that the B55α is required for dephosphorylation and activation of PDE4D5, which reduces cAMP/PKA signaling to promote NFκB activation in cells on FN.

### B55α directly binds N-terminal domain of PDE4D5

The identification of B55α as the regulatory/targeting subunit that mediates association of the PP2A complex with PDE4D5 at sites of adhesion raises the question, is this interaction direct? We therefore measured binding of purified recombinant B55α to purified recombinant fragments of PDE4D5 (Fig. 4A). B55α specifically bound to the N-terminal domain (amino acid 1-122) of PDE4D5 (Fig. 4B). Deleting this region from the S651E mutant of PDE4D5 (which binds PP2A more efficiently than WT; Fig 4C) abolished the interaction with B55α (Fig. 4C). Thus, the N-terminal domain of PDE4D5 mediates direct binding.

**Figure 4.**
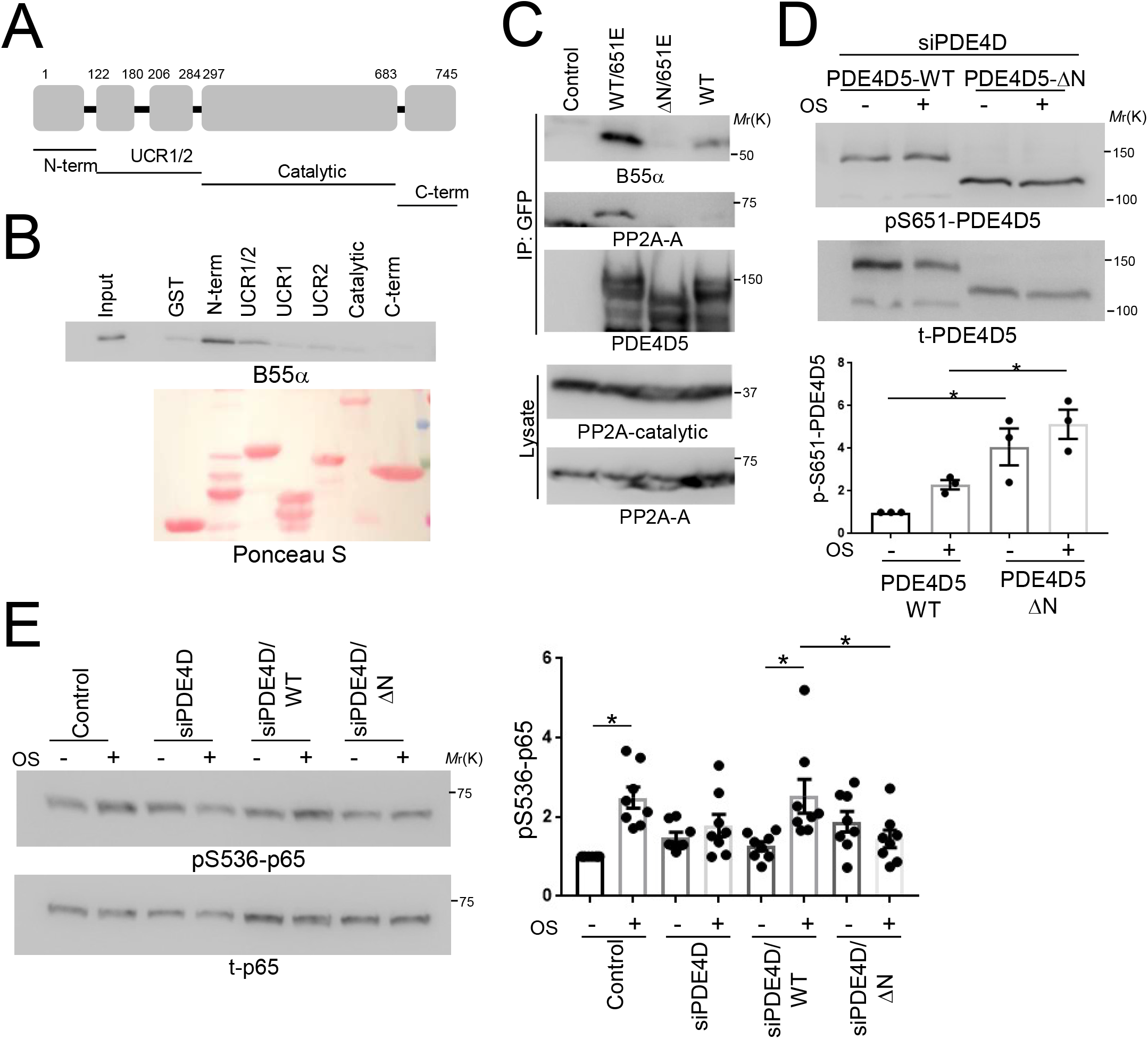
Mapping the B55α binding site on PDE4D5. **A.** Schematic representation of PDE4D5 fragments. **B.** Immobilized GST-tagged fragments of PDE4D5 were incubated with purified B55α, washed and probed for bound B55α (n=3). **C.** HEK 293T cells were transfected with the indicated GFP-tagged PDE4D constructs. Lysates were immunoprecipitated with GFP-Trap and immune complexes probed for B55α and PP2A-A subunit (n= 3). **D.** PDE4D5 knockdown BAECs were rescued with WT or N-terminally deleted mutant (DN). Cells were replated on FN and subject to OS for 2 hrs. PDE4D5 S^651^ phosphorylation was assayed as in Fig 3 (n=3). **E.** The same lysates were probed for p-p65 to assay NF-kB activity (n=8). *p<0.05 by one way ANOVA

To test the functional role of this interaction, we asked whether the N-terminal domain is required for PDE4D5 S651 dephosphorylation. Thus, endogenous PDE4D5 was knocked down and cells rescued with WT or the ΔN mutant. PDE4D5 S651 phosphorylation was increased with the ΔN mutant relative to WT protein in cells on FN (Fig. 4D). To test its role in inflammatory activation by disturbed flow, we carried out a similar knockdown-rescue protocol and then exposed the cells to oscillatory flow for 2h. As expected, NF-κB activation by disturbed flow was blocked by PDE4D5 depletion and rescued by WT PDE4D5 but not the ΔN mutant (Fig 4E). Together, these data show that direct binding of PDE4D to B55α promotes PDE4D5 dephosphorylation, which facilitates EC inflammatory activation by disturbed flow.

### FN promotes PP2A-B55α holoenzyme assembly

During these analyses, we unexpectedly observed that plating cells on FN greatly increased the co-immunoprecipitation between GFP-B55α and the catalytic subunit of PP2A compared to non-adherent cells (Fig 5A) or cells on MG (Fig. 5B). To confirm these results, we immunoprecipitated endogenous B55α, which also showed increased association with the catalytic subunit when plated on FN relative to MG (Fig. 5C). To test whether this effect requires PDE4D, we carried out similar co-IP experiments after knockdown of PDE4D. PDE4D5 depletion blocked assembly of the PP2A complex in cells on FN (Fig. 5D). Thus, FN promotes the assembly of a PP2A phosphatase complex, which requires PDE4D5. To our knowledge, this is the first instance where a receptor interaction controls assembly of a phosphatase complex.

**Figure 5.**
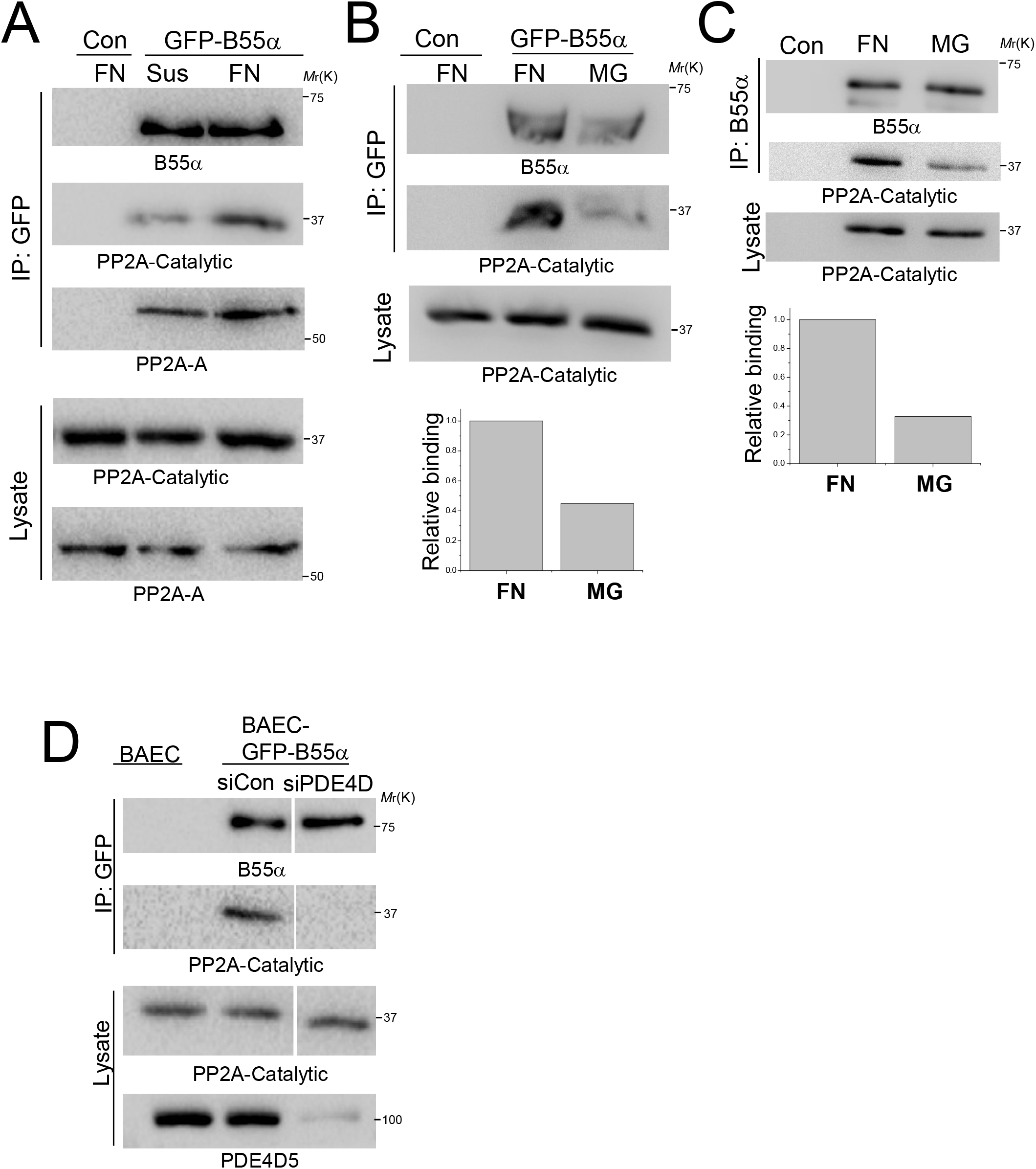
ECM-dependent PP2A-B55α assembly. **A.** BAECs expressing GFP-B55α were detached and replated on FN or kept in suspension for 20 min. GFP-B55α was immunoprecipitated and PP2A subunits were probed in the immune complexes. Sus; suspension **B.** BAECs expressing GFP-B55α were replated on FN or MG. GFP-B55α binding with B subunits was assayed as (A). (n=3) **C.** BAECs were replated on FN or MG. Endogenous B55α was immunoprecipitated and catalytic subunit assessed by Western blotting. **D.** BAECs transfected with PDE4D siRNA were replated on FN for 20 min. GFP-B55α was immunoprecipitated and catalytic subunit assayed by Western blotting as in (A) (n=3).

These results suggest that B55α may localize to cell-matrix adhesions through its association with PDE4D5. To test this idea, cells expressing GFP-B55α were plated on FN or on MG and examined with or without application of shear stress. Shear stress induced B55α relocalization to adhesions that partially overlapped paxillin (Fig. S4A, B). Based on the round shape of the adhesions, we examined whether those adhesions are Spreading Initiation Centers (SICs) that form transiently during cell spreading (35). RACK-1, a SIC marker, showed flow-dependent colocalization with B55α on FN (Fig. S4C). Shear stress induced B55α localization to sites of adhesion occurred preferentially on FN compared to MG (Fig. S4A). These results suggest that binding of cells to FN promotes PP2A holoenzyme assembly at sites of adhesion through an integrin-PDE4D5-B55α pathway.

### Yap dephosphorylation by PP2A-B55α

Cells contain tens of thousands of phosphorylated residues and many fewer phosphatase complexes; the finding that FN and integrin α5 control assembly of a major phosphatase complex therefore immediately suggests that there may be other functionally relevant substrates. The transcriptional regulator Yap is linked to integrin signaling, endothelial inflammation and atherosclerosis (36, 37). In ECs, disturbed flow induces Yap dephosphorylation, stabilization, nuclear translocation and transcription of target genes including inflammatory mediators (36, 37). Yap phosphorylation by the upstream kinase LATS1/2 inhibits its function via phosphorylation of S^127^, which results in Yap retention in the cytoplasm (38), and S^381^, which triggers Yap degradation (39). Yap may be dephosphorylated by PP2A in virally transformed cells (40). However the phosphatases that reverse these events under normal circumstances are poorly defined. We therefore examined Yap in our system.

To test whether Yap phosphorylation is matrix-dependent, ECs were plated on FN vs MG. Adherence to FN but not MG triggered rapid Yap S^127^ dephosphorylation (Fig 6A). Disturbed flow induced Yap phosphorylation in cells on MG, which was completely suppressed on FN (Fig 6B). These results indicate that a collagen/laminin basement membrane represses Yap activation, consistent with vascular stabilization, whereas FN promotes Yap activation, consistent with its pro-inflammatory effects.

**Figure 6.**
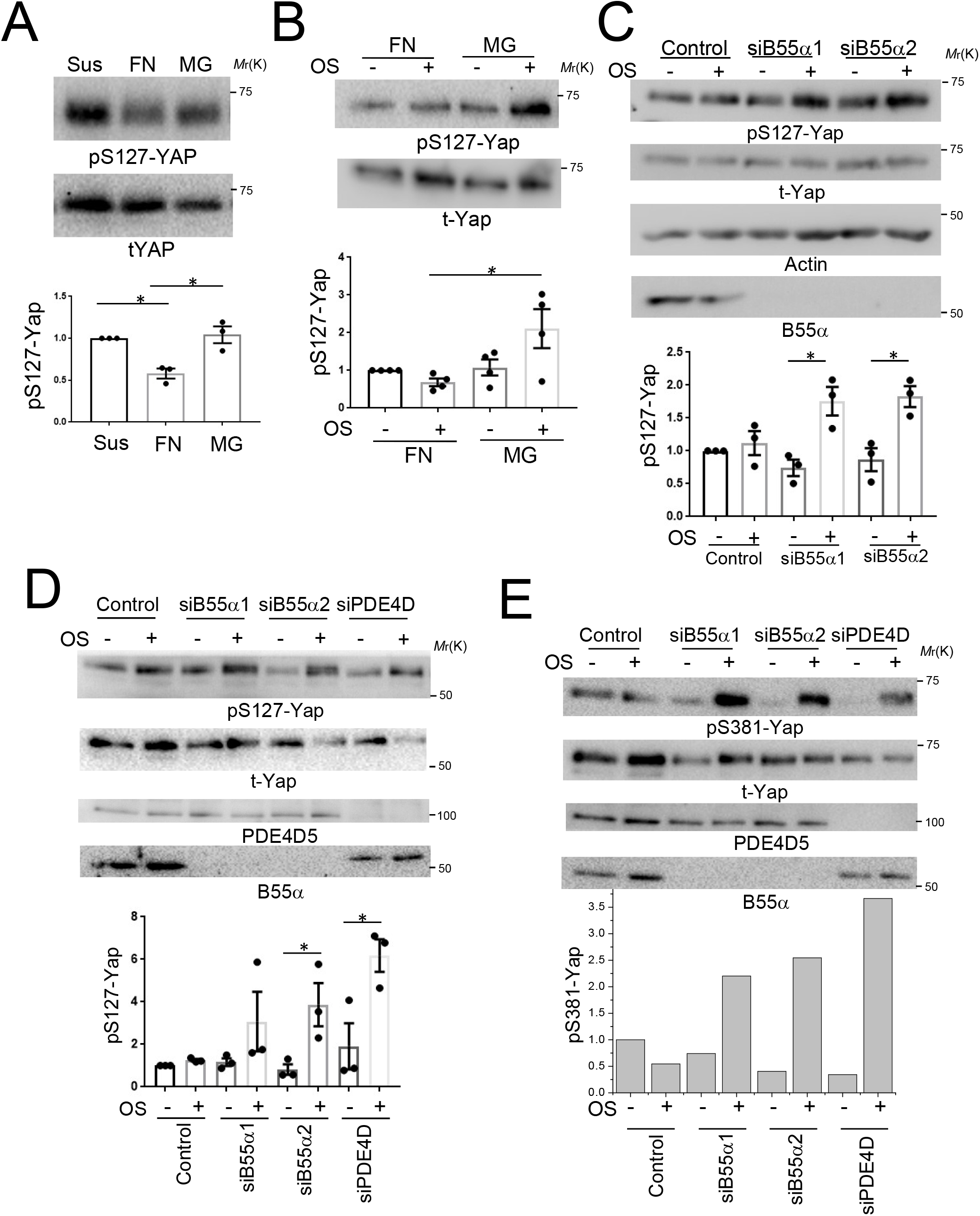
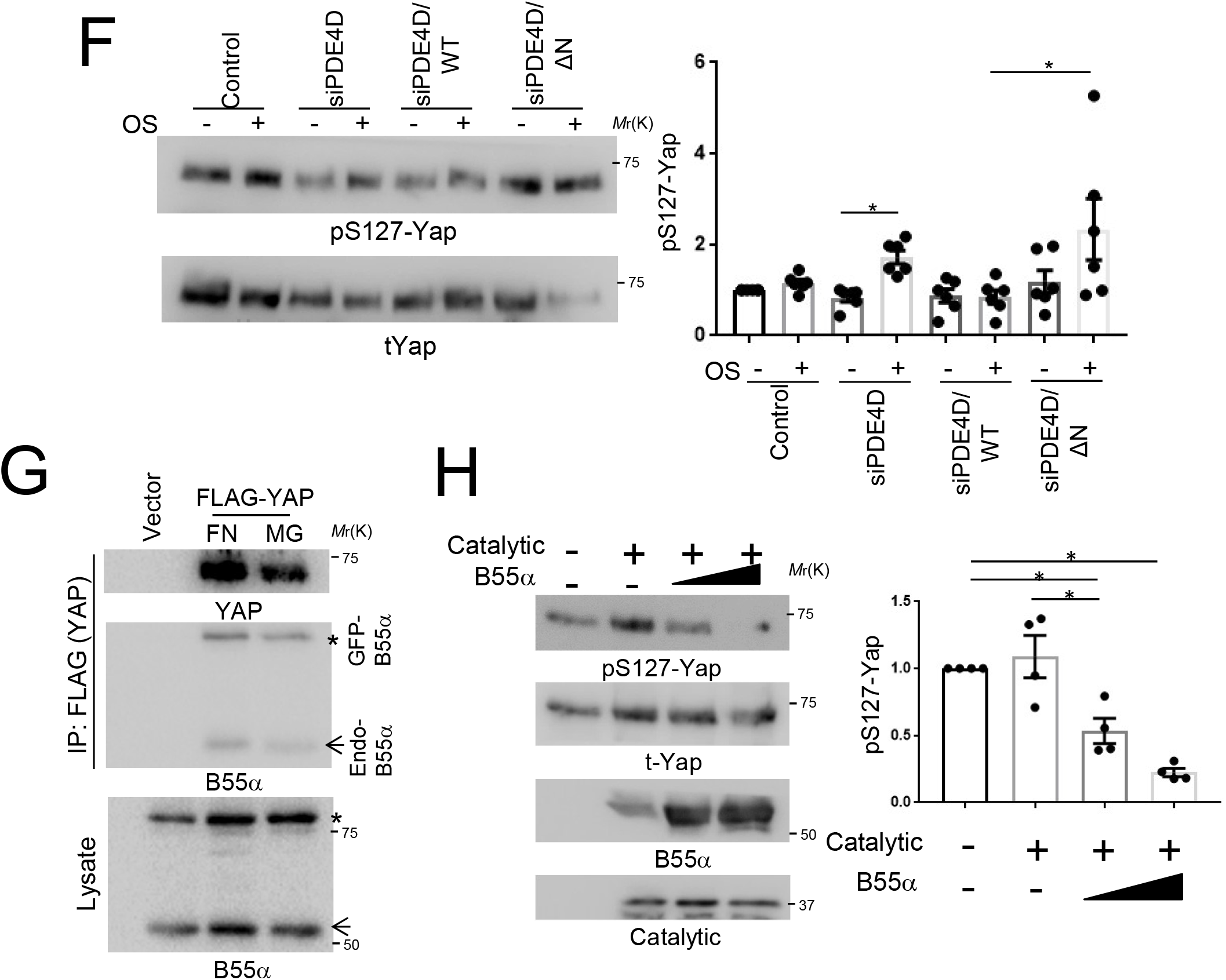
Yap is a PP2A-B55α substrate. **A.** BAECs were detached and kept in suspension (Sus) or replated on the indicated matrix protein for 1 hour. pS^127^-Yap was measured by immunoblotting (n=3). **B.** BAECs plated on FN or MG for 6 hr were exposed to OS for 2 hr. Yap S^127^ phosphorylation was assayed as in A (n=4). **C.** BAECs transfected with B55α siRNA were detached and replated on FN, then exposed to OS for 2 hr. Yap phosphorylation was assayed as in A (n=3). **D, E.** HUVECs transfected with B55α siRNA or PDE4D siRNA were detached and replated on FN, then sheared for 2 hr. Lysates were analyzed by Western blotting for pS^127^-Yap (D) or pS^381^-Yap (E). n=3 independent experiments. **F.** PDE4D5 knockdown BAECs were reconstituted with WT or N-terminally deleted mutant (DN). Cells were replated on FN, subject to OSS for 2 h, and Yap S^127^phosphorylation assayed as in A (n=6). **G.** BAECs expressing GFP-B55α were transfected with FLAG-Yap and replated on FN or MG for 20 min. Yap was immunoprecipitated with FLAG antibody and probed for B55α. N= 3 independent experiments. **H.** In vitro phosphatase assay. PP2A catalytic subunit was immunoprecipitated with PP2A C subunit antibody (clone 1D6) and incubated for 30 min at 37°C with phosphorylated Yap in the presence of increasing amount of purified B55α. Reaction mixtures were immunoblotted for pS^127^-Yap (n=4). *p<0.05 by two-tailed t-test (C, D, F) or one way ANOVA (A, B, H)

To test whether these matrix-specific effects involved the PP2A complex, cells were transfected with B55α siRNA. Yap dephosphorylation in cells on FN was prevented by knock-down of B55α (Fig. 6C-E). Yap dephosphorylation was also blocked by knockdown of PDE4D (Fig. 6D-E), which was rescued by WT but not the B55α-binding deficient PDE4D-ΔN mutant (Fig. 6F). Yap nuclear translocation also increased on FN (Fig S5A), which was blocked by knockdown of B55α or PDE4D (Fig S5B). Yap S^127^ dephosphorylation correlated with induction of Yap target genes, ANKRD1 and Cyr61, which was also blocked by B55α or PDE4D knock-down (Fig. S5C). Consistent with reduced Yap S^381^ dephosphorylation, the previously reported increase in Yap protein levels under longer term disturbed flow(36) was abolished by knockdown of B55α or PDE4D (Fig S5D). Together, these results show that the integrin-PDE4D5-B55α axis controls Yap activity.

To identify the direct PP2A-B55α target in the Hippo-Yap pathway, we first examined the upstream kinase LATS1. In cells on FN, flow decreased LATS1 phosphorylation, however, this was unaffected by B55α knock-down (Fig. S6). This result excludes LATS1 and all upstream steps in the pathway. We therefore examined Yap itself. Immunoprecipitated Flag-Yap associated with both GFP-B55α and endogenous B55α (Fig. 6G). To test whether Yap is a direct PP2A substrate, we measured Yap dephosphorylation in vitro with purified components. Highly phosphorylated Yap purified from confluent 293T cells was incubated with PP2A catalytic subunit immunoprecipitated from endothelial cells using an antibody (clone 1D6) that specifically recognizes catalytic subunit without bound B subunit (41). Catalytic subunit alone showed negligible activity toward phospho-Yap, however, addition of purified B55α induced dose-dependent Yap S^127^ dephosphorylation (Fig. 6H). Thus, pYap is a direct, B55α-dependent PP2A target.

These results predict that the previously reported activation and stabilization of Yap in regions of disturbed flow in vivo (36, 37) should require the FN-integrin-PDE4D5-B55α pathway. To test this hypothesis, we compared vessels from WT versus PDE4D^mut^ mice on the ApoE^−/−^ background on a chow diet, which have increased vascular inflammation but minimal plaque formation. The increased Yap accumulation in ECs in the atheroprone inner curvature of the aortic arch from WT mice was significantly blunted in PDE4D^mut^ mice (Fig. 7A). Furthermore, expression of Yap target gene, Cyr61 and CTGF in the inner curvature was significantly decreased in PDE4D^mt^ mice (Fig. 7B, C). Together, these data identify Yap as a direct target of the PP2A-B55α complex that is regulated through PDE4D5.

**Figure 7.**
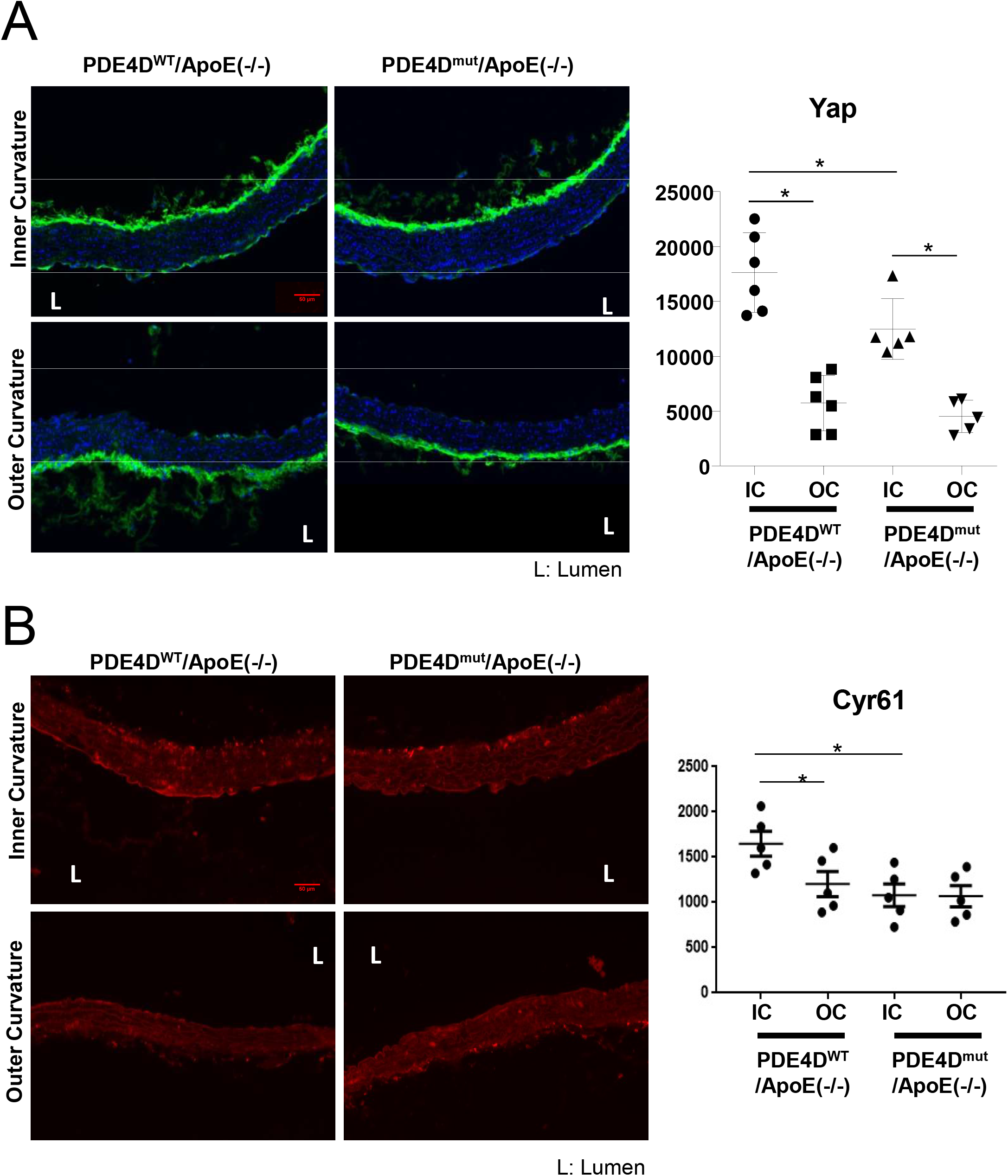
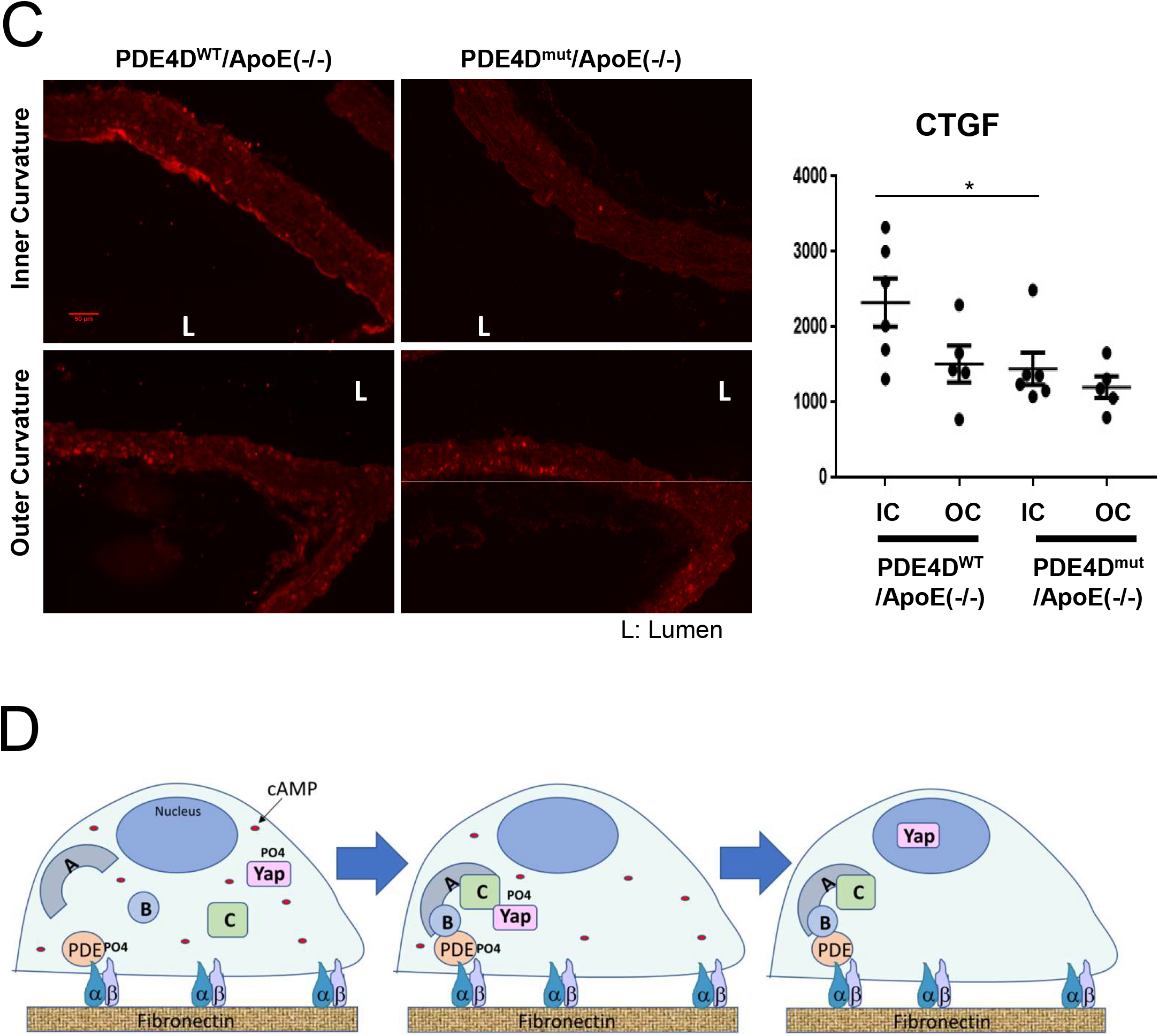
Effect of PDE4D mutation on Yap and target gene expression in disturbed flow region. **A-C**. Aortas from PDE4D^WT^/ApoE null mice or PDE4D^mut^/ApoE on normal chow were fixed, longitudinally sectioned and stained for DAPI to identify nuclei and for total Yap (A). The adjacent sections were stained for Yap target genes, Cyr61 (B) and CTGF (C). Mean intensity in the endothelial layer of the inner curvature (IC) and outer curvature (OC) was quantified (arbitrary units). *p<0.05 by two-tailed t-test. **D.** Model for PDE4D-dependent regulation of PP2A and its substrate Yap.

## Discussion

These results show, first, that blocking the binding of PDE4D5 to integrin α5 reduced EC inflammatory activation and atherosclerotic plaque burden, and increased plaque stability. Together with the previous findings that mutation of the integrin α5 cytoplasmic domain inhibited these events, these data provide conclusive evidence that the α5-PDE4D5 interaction contributes to inflammatory activation of the endothelium and to atherosclerosis. The evidence of increased plaque stability is notable in light of the decreased plaque stability after deletion of plasma FN (26). FN therefore appears to promote inflammation and plaque expansion through α5/PDE signaling, whereas the enhancement of plaque stability is separable, probably through FN’s role in scaffolding collagen fibrillogenesis (42). The PDE4D5 mutation also slightly improved, or at least did not inhibit, vascular remodeling in the hindlimb ischemia model. The integrin-PDE4D5 pathway, unlike FN itself, may therefore be a suitable therapeutic target.

Investigating the mechanism of PDE4D5 activation led to the unanticipated finding that FN and PDE4D5 control assembly of a B55α-containing PP2A complex. This complex localizes to sites of adhesion and dephosphorylates the inhibitory S^651^ site on PDE4D5. PDE4D5 is therefore both upstream and downstream of PP2A. Given the paucity of phosphatase complexes compared to phosphorylated residues in the proteome, other relevant B55α-PP2A substrates seemed likely. As large scale phospho-proteomic profiling would not distinguish direct B55α-PP2A substrates from secondary changes, we took a candidate approach. Several literature reports suggested the Hippo-Yap pathway as a possible target. Indeed, plating cells on FN induced Yap dephosphorylation on key regulatory sites, indicative of Yap activation. Adherence to FN also blocked disturbed-flow induced Yap phosphorylation (inhibition), increasing nuclear translocation and Yap-target gene expression, and stabilizing Yap protein. The B55α-PP2A complex directly dephosphorylates Yap, as indicated by lack of effect on the upstream kinase LATS, co-IP of Yap and PP2A, and in vitro dephosphorylation with purified components. Consistent with these in vitro results, the reported stabilization of Yap at atheroprone regions of arteries (36, 37) was blunted in PDE4D5^mut^ mice. As Yap inactivation in ECs potently reduced atherosclerosis in mice (36, 37), this pathway appears to substantially contribute to the effects of the FN-α5-PDE4D5 pathway in vascular disease.

These experiments were done using matrix proteins adsorbed at low concentrations to rigid glass or plastic, thus, are distinct from the regulation of Yap by matrix stiffness (43, 44). The results, however, fit with the well-known ability of FN to promote EC cell cycle progression relative to laminin and collagen (45), as well as inflammatory activation (2, 9). Both mechanical and biochemical signaling mechanisms may contribute in vivo, where atherosclerosis and aging are associated with increased matrix stiffness (46). How specific integrin α5 signaling and matrix stiffness interact to regulate Yap activation and gene expression will be interesting to explore in future work.

Binding of cell-derived FN that contains the alternatively spiced EDA domain, but not plasma FN, to TLR4 has been implicated in flow-dependent vascular inflammation and atherosclerosis (16, 47). Supporting this idea, both TLR4 and EDA-positive FN are increased in atherosclerotic lesions (48). Interestingly, there is considerable overlap between the effects of EDA deletion in these studies and blockade of PDE4D binding in our work. This could be explained if TLR4 pro-inflammatory signaling is dampened by cAMP/PKA, such that EDA-TLR4 signaling requires the integrin α5-PDE4D pathway. EDA also promotes FN matrix assembly (49), thus, deletion of the EDA domain may decrease the amount of FN in the vessel wall, thus influencing all FN-dependent pathways.

The molecular mechanism by which PDE4D5 controls B55α-PP2A assembly is unknown. In vitro Yap dephosphorylation experiments indicate that PDE4D5 is not required for the B55α-PP2A assembly in solution, as expected from published data (50). PDE4D5 might block inhibitory factors or protein modifications of B55α to facilitate holoenzyme assembly in vivo. Elucidation of detailed mechanism is an important question for further analysis.

Atherogenesis shares features with the physiological vascular remodeling that promotes efficient perfusion of tissues via angiogenesis and modulation of vessel diameter (10, 11). These processes involve changes in the extracellular matrix (ECM), inflammatory activation of the endothelium and recruitment of leukocytes that assist with remodeling, followed by resolution of inflammation and restoration of stable vasculature. EC responses to disturbed flow at artery branches and regions of high curvature may be seen as futile, flow-dependent remodeling that synergizes with systemic inflammatory and metabolic stresses to induce atherogenesis. By contrast, physiological remodeling is suppressed by elevated inflammatory status and metabolic disorders (51–54). It is well-accepted that cAMP/PKA activation is anti-inflammatory and reduces a plaque burden (55–57). However, cAMP/PKA signaling is highly multifunctional with distinct effects in the heart, brain, liver and other tissues, thus, is not an attractive target for systemic therapy. Solving vascular disease requires more precise methods to reduce chronic vascular inflammation without harmful side effects. Interactions within the α5-PDE4D5-PP2A pathway therefore appear to be promising therapeutic targets.

In summary, our data elucidate a pathway in which FN binding to the integrin α5 subunit recruits PDE4D5 to induce assembly of a PP2A-B55α complex that regulates flow-induced Yap signaling and very likely other functionally relevant effectors. This pathway is likely to have widespread implications for control of cell phenotype by ECM across a range of systems. Important questions for future work include discovering the mechanism by which PDE4D enhances PP2A-B55α association on FN, identifying additional matrix-dependent substrates for PP2A, and elucidating their roles in EC phenotype and vascular disease.

## Methods

### Generation of PDE4D^mut^ mice

We used the double nicking strategy to minimize off-target effects (24, 58, 59). Two guide RNA sequences were selected from candidate sequences using algorithm from http://crispr.mit.edu/. Guide A: CUUUUCCUUCUGAGUUGGAGAGG, Guide B: GGCCGAUGUCGCAGAUCAGUGGG. The donor ssDNA oligonucleotide (ssODN) template including PDE4D mutations was synthesized (IDT; Ultramer grade) and the sequence follows 5’-atgagtcggtctggcaaccaggtgtcggagtacatctcaaacacatttctcgataagcaacatgaagtggaaatcccctctccaacgcagaaggaaaaagaggaggaggaagagccgatgtcgcagatcagtggggtcaagaagttgatgcacagctccagcctaactaattcatgtatc-3’ Templates for in vitro transcription of the sgRNAs were PCR amplified from the pX335 plasmid (a gift from Feng Zhang, Addgene plasmid # 42335) using the following primers: Forward primer for Guide A: 5’tgtaatacgactcactataggCTTTTCCTTCTGAGTTGGAGgttttagagctagaaatagc 3’ (T7 promoter/guide/sg scaffold template) Forward primer for Guide B: 5’ tgtaatacgactcactataggGGCCGATGTCGCAGATCAGTgttttagagctagaaatagc 3’ Reverse primer: 5’ AGCACCGACTCGGTGCCACT 3’

The DNA template for in vitro transcription of Cas9 mRNA was PCR amplified from pX335, in vitro transcribed and purified as described (58). Cas9 mRNA (100 ng/μl), guide A and guide B sgRNAs (50 ng/μl each), and the ssODN template (100 ng/μl) were microinjected into the cytoplasm of C57Bl/6J zygotes; surviving zygotes were transferred to oviducts of CD-1 pseudopregnant females the day of injection as described (60).

For genotyping, the following primers were used: Wild type allele: 200 bp PCR product, forward primer: 5’gataccaacttgtagtcaactc3’, reverse primer: 5’tcggccttttcttcttctcctt3’.

Mutant allele: 200 bp PCR product, same forward primer as wild type allele, reverse primer: 5’tcggctcttcctcctcctcttt3’.

### Histology and immunohistochemistry

Mice were euthanized according to the Yale University IACUC protocol and perfused via the left ventricle with PBS and then 3.7 % formaldehyde for fixation. For immunohistochemsitry, longitudinal paraffin sections of aortas were prepared by Yale Pathology Tissue Services. For cryosections of aortic roots, fixed tissue was kept in 30% sucrose in PBS at 4 °C overnight, incubated with a 1:1 mixture of OCT and 30% sucrose/PBS for 30 minutes and embedded in OCT on dry ice. Aortic root cryosections (10 µm) were prepared with a cryostat and incubated in blocking buffer containing 5% goat serum/0.1 % TX-100/PBS 1 hour at RT. Sections were incubated in primary antibodies at the indicated concentrations in blocking buffer overnight at 4°C, washed 3 times in PBS, then incubated with Alexa Flour 568-conjugated secondary antibodies (1:300, Invitrogen) 1 hour at RT. After washing with PBS, sections were mounted in Fluoromount G (Southern Biotech). Antibodies used were anti-VCAM1 (Abcam, ab955, 1/300), anti-ICAM1 (Biolegend, 116102, 1/400), anti-p-NFκB (Abcam, ab28856, 1/400), anti-CD68 (Abcam, ab955, 1/400), anti-smooth muscle actin (Sigma, C6198, 1/500), anti-MMP2 (SantaCruz, sc10736, 1/100), anti-MMP-9 (Abcam, ab38898, 1/500). Six mice (3 male and 3 female, 3 month old) were examined for inflammatory markers in the aortic arch for each strain. For atherosclerosis, 3 month old homozygous PDE4D^mut^;ApoE^−/−^ mice were placed on a high fat diet (Open Source, D12079B) for 4 months, then euthanized, fixed and examined as described above. For histology, Oil Red O (0.3%, Sigma-Aldrich) and Picrosirius Red stain (Abcam) were used. All stained sections were imaged on a Nikon 80i microscope. Image analysis was done using Image J software. Positively stained areas of Oil Red O–stained aortic roots were quantified and expressed as the percentage of the total cross-section area of aortic root. Sirius Red positive areas were expressed as the percentage of plaque area.

### Hindlimb Ischemia

Femoral arteries were ligated at two locations spaced 5 mm apart and the arterial segment between the ligation was excised. Blood flow in the hindlimbs was imaged using a Moor Infrared Laser Doppler Imager (LDI; Moor Instruments Ltd) at 37.0°C to 38.0°C under ketamine/xylazine (80/5 mg/kg) anesthesia. Data were analyzed with Moor LDI image processing software V3.09 and plotted as the ratio of flow in the right/left (R/L) hindlimb.

### Micro-CT Angiography

2D mCT scans were performed with a GE eXplore MS Micro-CT System, using a 400 cone-beam with angular increment of 0.5 degrees and 8 to 27 μm slice thickness at a voltage of 163.2 mAs, 80 kVp. Data were transferred to a Dell Dimension computer with 3D volume rendering software (version 3.1, Vital Images Inc, Plymouth, MN) and microview software (version 1.15, GE medical system). ImageJ and Image Pro Plus (Media Cybernatics) software were used to analyze vessel number, diameter, area, volume, and arterial density.

### In situ zymography

Unfixed aortic root cryosections were incubated with DQ-gelatin (20 μg/ml) in gelatinase reaction buffer at 37□ for 1 hour, washed with PBS and mounted with antifade mounting solution.

### Cell culture

BAECs were grown in DMEM containing 10% FBS and penicillin/streptomycin. HUVECs were grown in M199, 20% FBS, 5 mg/ml ECGS, 100 μg/ml heparin, penicillin/streptomycin.

### Shear stress experiments

Serum-starved endothelial cells were replated on glass slides coated with the indicated proteins for 6 hr before application of flow. The slides were loaded into parallel plate flow chambers. Laminar shear at 15 dynes/cm^2^ was used to mimic high flow in athero-resistant regions of arteries. Oscillatory shear of 1 ± 5 dynes/cm^2^, 1 Hz was used to mimic disturbed flow in athero-prone regions.

### Immunoprecipitation

Cells were lysed in 20 mM PIPES pH 6.8, 1% TX-100, 150 mM NaCl, 150 mM sucrose, 0.2% sodium deoxycholate, 500 μM EDTA and protease inhibitors. After incubation on ice for 15 min, samples were centrifuged at 20,000g for 10 min, supernatants were diluted 10X in buffer containing 20 mM PIPES pH 6.8, 1% TX-100, 150 mM NaCl, 150 mM sucrose, 2.5 mM MgCl_2_ and 2.5 mM MnCl_2_. For immunoprecipitation of GFP-B55α, GFP trap agarose beads (ChromoTek) were incubated with the lysates for 3 hr at 4 °C before washing with dilution buffer. For immunoprecipitation of endogenous B55α, B55α antibody (clone 2G9, Milipore) was covalently cross-linked to agarose beads using Pierce Crssolink IP kit (Thermo Fisher Scientific).

### Immunoblotting

Antibodies used for Western blotting are following.

α-p-NFκB p65 (S536): rabbit mAb (93H1), Cell Signaling (3033L), 1/1,000.

α-NFkB-p65: mouse mAb (F-6), Santa Cruz (sc-8008), 1/1,000.

α-PP2A, C subunit: mouse mAb, BD Transduction Laboratories (610556), 1/1,000.

α-PP2A, B55α: mouse mAb (2G9), Cell Signaling (5689s), 1/1,000.

α-PP2A, A subunit: rabbit mAb (81G5), Cell Signaling (2041s), 1/1,000.

α-p-LATS1(T1079): rabbit mAb (D57D3), Cell Signaling (8654s), 1/1,000.

α-LATS1: rabbit pAb, Abcam (ab70565), 1/1,000.

α-p-Yap (S127): rabbit pAb, Cell Signaling (4911s), 1/1,000.

α-p-Yap (S381): rabbit mAb (D1E7Y), Cell Signaling (13169s), 1/1,000.

α-Yap: mouse mAb (63.7), Santa Cruz (sc-101199), 1/200.

α-p-eNOS (S635): rabbit pAb, Upstate (07–562), 1/1,000.

α-eNOS: rabbit pAb, BD (610298), 1/1,000

Band intensities from immunoblotting were quantified by densitometry using imageJ software.

### GST pull-down assays

5 μg of GST fusion proteins on GSH-agarose beads were incubated with 1.5 μg of B55α purified from insect cells in buffer containing 20 mM PIPES pH 6.8, 1% TX-100, 500 mM NaCl, 0.1% sodium deoxycholate, 150 mM sucrose and 1 mg/ml BSA for 30 min at 4 °C, then washed and analyzed by SDS-PAGE and Western blotting.

### Plasmids and siRNA

Human PDE4D5 wild type and PDE4D5-dN mutant were PCR amplified and cloned into pLVX-mCherry-N1 vector (Clontech) using Gibson assembly after vector digestion with XhoI and EcoRI. WT forward primer; 5’-agcgctaccggactcagatctcgagatggctcagcagacaagcccgg, dN forward primer; 5’-agcgctaccggactcagatctcgaatgcaacgacgggagtccttcctgtatc, reverse primer; 5’-cgcggtaccgtcgactgcagaattcgcgtgtcaggagaacgatcatcta. Human B55α was PCR amplified from HUVEC cDNA and cloned into pEGFP-C1 (Clontech) using HindIII and BamHI sites. Forward primer; 5’-gcaagcttcgatgttcccgaagttttctcttcg, reverse primer; 5’-gcggatccctaattcactttgtcttgaaatat. GFP-tagged B55α was PCR amplified from pEGFP-C1-B55α and cloned into pLPCX (Clontech) using NotI and ClaI sites. Forward primer; 5’-gcgcggccgcatggtgagcaagggcgagga, reverse primer; 5’-gcatcgatctaattcactttgtcttgaaatat. The siRNA sequence for PDE4D used in BAECs and HUVECs was 5’-AAGAACUUGCCUUGAUGUACA-3’ (9). The siRNA for B55α were #1, 5’-GCAGAUGAUUUGCGGAUUAUU-3’, #2, 5’-GUGCAAGUGGCAAGCGAAAUU-3’. PDE4D5 fragments cloned into pGEX-4T1 were previously described (61).

### Proteomic analysis for PDE4D5 binding proteins

GFP-tagged PDE4D5-S651E mutant was stably expressed in BAECs using retroviral infection. The cells were detached and kept in suspension for 30 min, replating on FN or matrigel for 20 min, lysed and immunoprecipited with GFP-Trap. After SDS PAGE and silver staining, specific bands were excised and submitted to Yale Keck Biotechnology Resource Laboratory for LC-MS/MS analysis.

### Statistics

Statistics were analyzed using Student’s t-test or 1-way ANOVA (for multiple comparisons) in GraphPad Prism 6. Statistical significance was taken as p<0.05. Data are represented as means ± SEM.

## Author contributions

SY and MAS designed the project. SY, RH, MES, ANS and ZZ conducted the experiments and analyzed the data. SY and MAS generated the figures and wrote the manuscript. AJK and DCP provided technical advice and edited manuscript. MAS directed and supervised the project.

## Acknowledgements

We thank Dr. Stefano Piccolo (University of Padova, Italy) for kindly providing FLAG-Yap DNA and for critical comments on the manuscript. Lipid analysis was done by the Yale Mouse Phenotypic Center, supported by grant U24 DK059635. This work was funded by National Institutes of Health grants 5R01HL75092, RO1 HL135582 and PO1107205 to M.A.S. A.J.K is funded by NIH funding R01s MH115939, NS105640, and NS089662. We are grateful to Rita Webber, Nicole Copeland and Laran Coon for maintaining mouse colonies used in this study.

