## Supplementary figures and legends for "Fibronectin receptor integrin α5β1 regulates assembly of PP2A complexes through PDE4D: modulation of vascular inflammation and atherosclerosis"

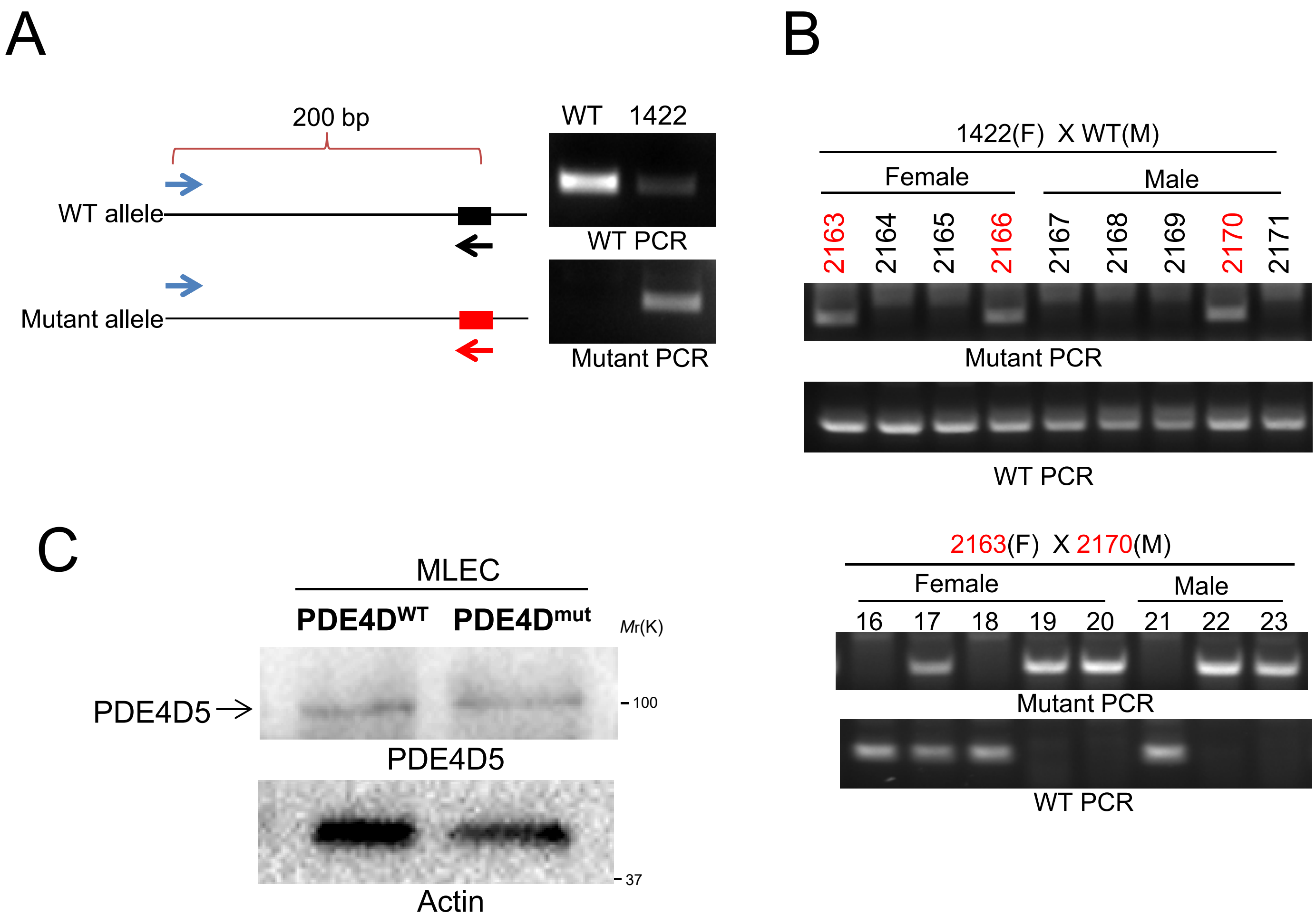


1. Generation of PDE4D$^{mut}$ mice

**A**. Genotype was screened by PCR using primers that discriminate between wild type and mutant alleles. **B**. Germ-line transmission. The mutant allele was successfully inherited in F1 and F2 offspring, including homozygous F2 pups. **C.** PDE4D5 protein levels. Endothelial cells from each strain were lysed and immunoblotted for PDE4D5.

Supplementary Fig. 1

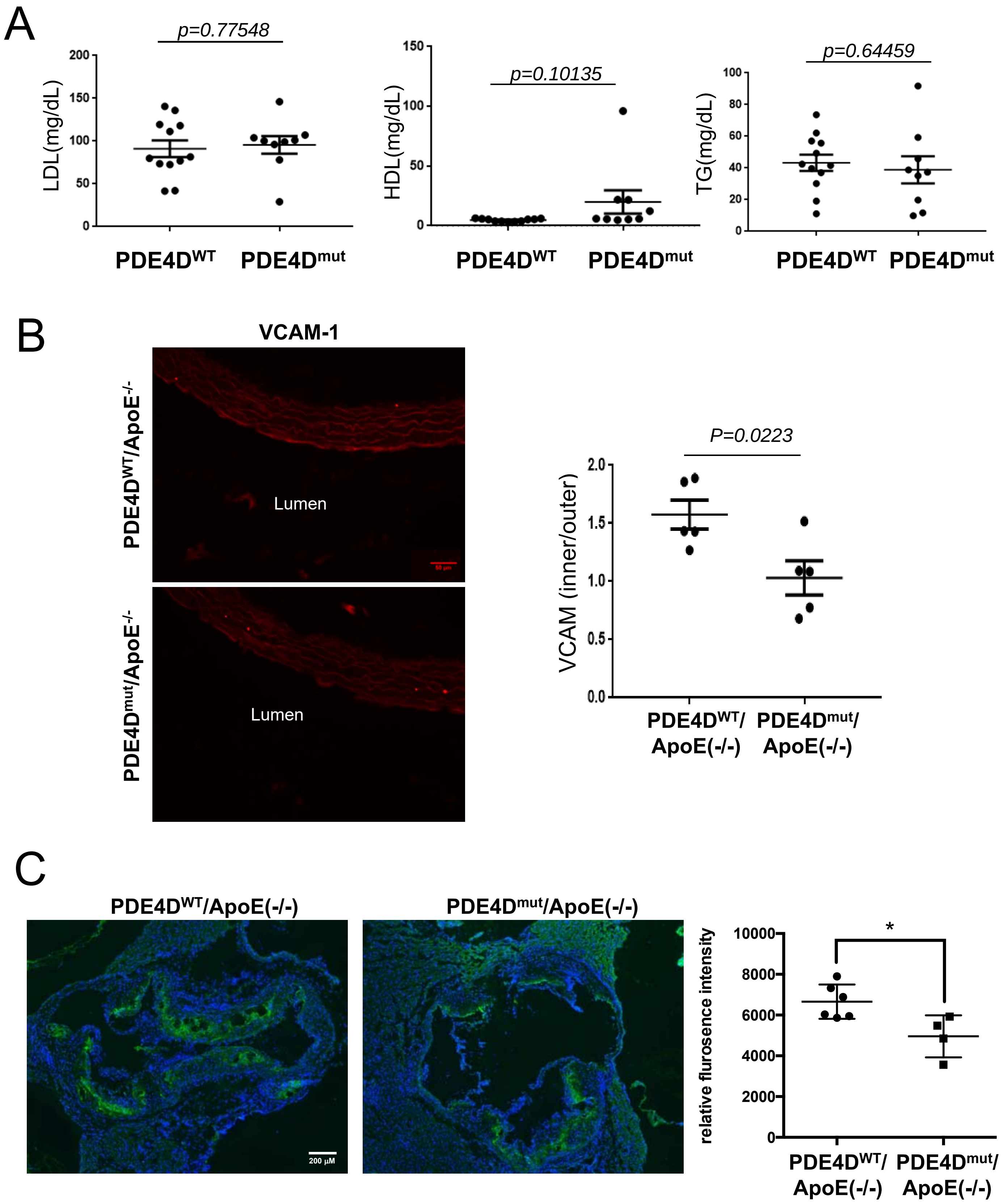


Supplementary Fig. 2

D

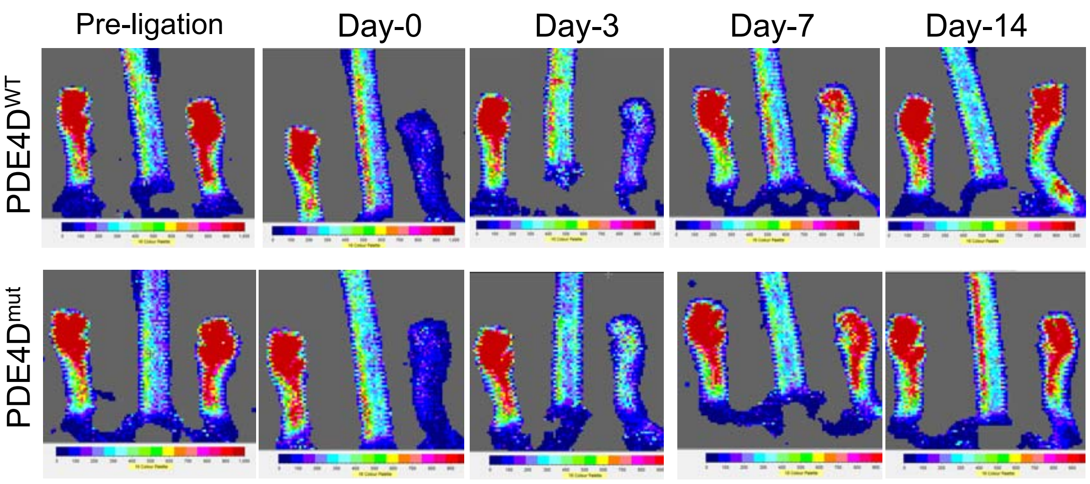


E

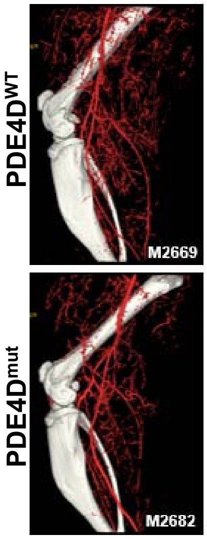


F

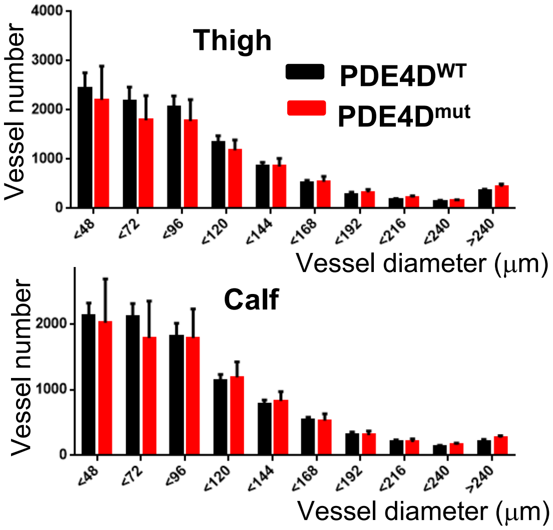


2. Characterization of PDE4D$^{mut}$ mice

**A**. Serum lipid profiles of PDE4D$^{mut}$; ApoE$^{-/-}$ mice on high fat diets (n=9-12). **B**. Longitudinal sections of aortas from WT or PDE4D$^{mut}$ ApoE null mice after 4 months on a high fat diet were stained for VCAM-1. Images show the inner curvature of the aortic arch. **C**. MMP activity within the plaque was measured using in situ zymography. **D**. Representative Doppler images of hindlimb blood flow at the indicated times after femoral artery ligation. **E, F**. Vessel density in the control leg 14 days after surgery by MicroCT. Results were quantified according to vessel diameter. *$p<0.05$ by two-tailed t-test

Supplementary Fig. 2

### LCMS Peptides

**Protein ID** 2ABA_PIG

**Protein Name** Serine/threonine-protein phosphatase 2A 55 kDa regulatory subunit B alpha isoform (Fragment) OS=Sus scrofa GN=PPP2R2A PE=2 SV=1

**Percent Coverage** 22.8

**5 peptides identified with score greater than identity score**

| Score | Expectation | Peptide Sequence | Start | End | M/Z | Ion Mass | Ion Mass(calc) | Delta | ppm | Charge |
|---|---|---|---|---|---|---|---|---|---|---|
| 86.58 | 1.1E-7 | R.VVIFQQEQENK.I | 31 | 41 | 681.3606 | 1360.7066 | 1360.6987 | 0.008 | 5.9 | 2 |
| 77.63 | 6.8E-7 | R.SFFSEIISSISDVK.F | 258 | 271 | 779.9067 | 1557.7989 | 1557.7926 | 0.0062 | 4 | 2 |
| 70.9 | 0.0000039 | K.NAAQFLLSTNDK.T | 85 | 96 | 661.3403 | 1320.666 | 1320.6674 | -0.0014 | -1.1 | 2 |
| 51.35 | 0.00034 | R.DITLEASR.E | 355 | 362 | 452.741 | 903.4674 | 903.4661 | 0.0013 | 1.4 | 2 |
| 41.53 | 0.0023 | R.INLWHLEITDR.S | 179 | 189 | 470.5894 | 1408.7462 | 1408.7463 | -0.0001 | -0.1 | 3 |

Export options: CSV | Excel

**One peptide identified with score between homology score and identity score**

| Score | Expectation | Peptide Sequence | Start | End | M/Z | Ion Mass | Ion Mass(calc) | Delta | ppm | Charge |
|---|---|---|---|---|---|---|---|---|---|---|
| 15.85 | 0.41 | R.GEYNVYSTFQSHEPEFDYLK.S | 48 | 67 | 818.3667 | 2452.0782 | 2452.0859 | -0.0077 | -3.1 | 3 |

Export options: CSV | Excel

**2 peptides identified with score less than homology score**

| Score | Expectation | Peptide Sequence | Start | End | M/Z | Ion Mass | Ion Mass(calc) | Delta | ppm | Charge |
|---|---|---|---|---|---|---|---|---|---|---|
| 8.63 | 6.6 | R.DKRPEGYNLK.E | 107 | 116 | 407.2202 | 1218.6389 | 1218.6357 | 0.0032 | 2.6 | 3 |
| 6.53 | 6.2 | R.PMDLMVEASPR.R | 138 | 148 | 623.3024 | 1244.5903 | 1244.5893 | 0.001 | 0.8 | 2 |

Export options: CSV | Excel

3. Liquid chromatography-mass spec identification of B55α

The fibronectin-specific 55 kDa band from Fig 3A was submitted for mass analysis. Multiple peptides identify it as the PP2A B55α subunit.

Supplementary Fig. 3

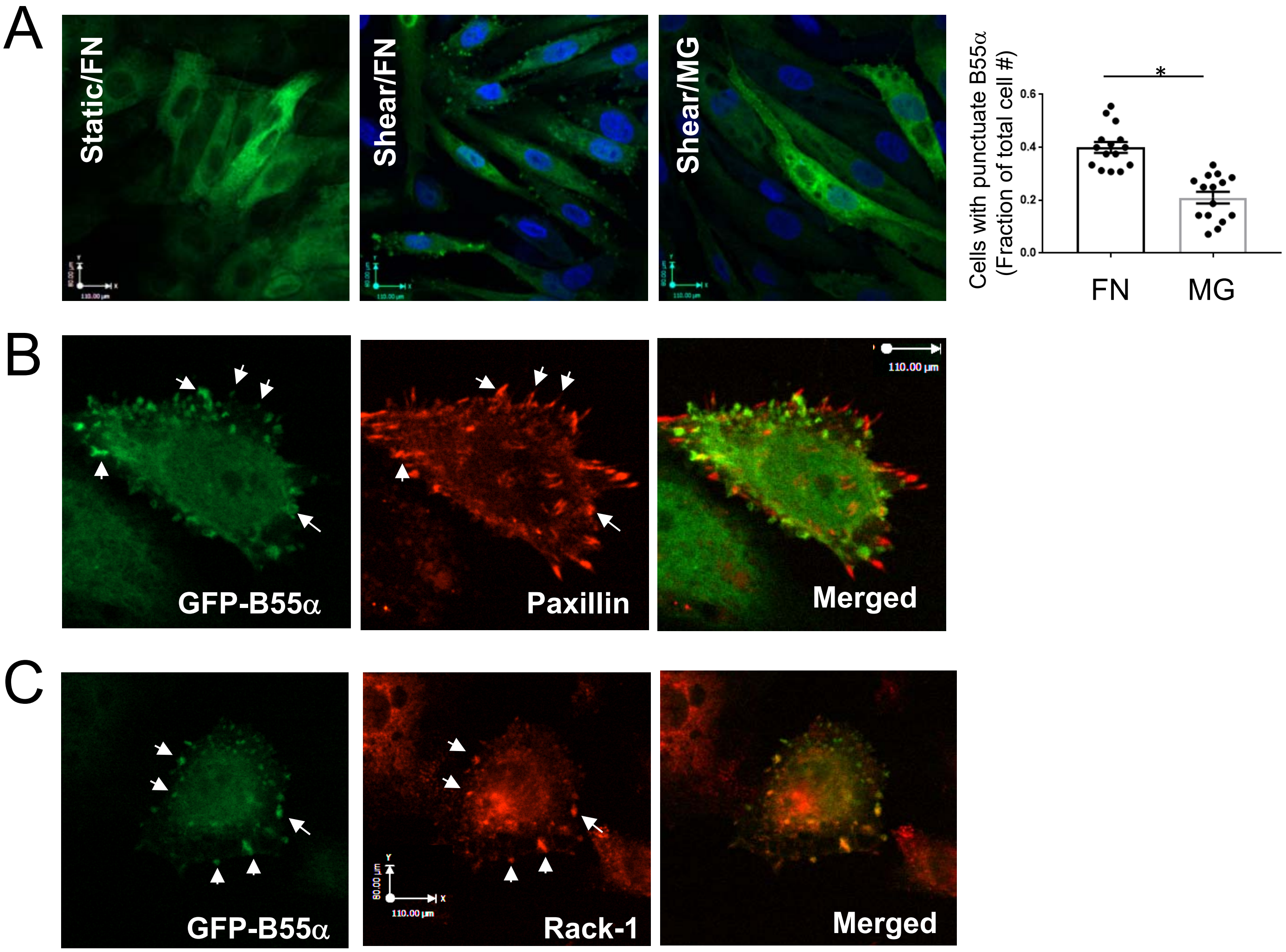


<u>4. ECM-dependent PP2A-B55α localization</u>

**A**. B55α localization. BAECs expressing GFP-B55α were plated on FN or MG and exposed to laminar shear for 30 min. After fixation, GFP fluorescence was imaged using confocal microscopy. The number of cells with punctuate B55α was compared between FN and MG (n=15 images pooled across three independent experiments). *p<0.05 by two-tailed t-test. **B**. BAECs expressing GFP-B55α and RFP-paxillin were plated on FN and subject to shear for 30 min. **C**. BAECs expressing GFP-B55α were plated on FN and exposed to shear for 30 min. After fixation, the cells were stained for RACK-1 and imaged.

Supplementary Fig. 4

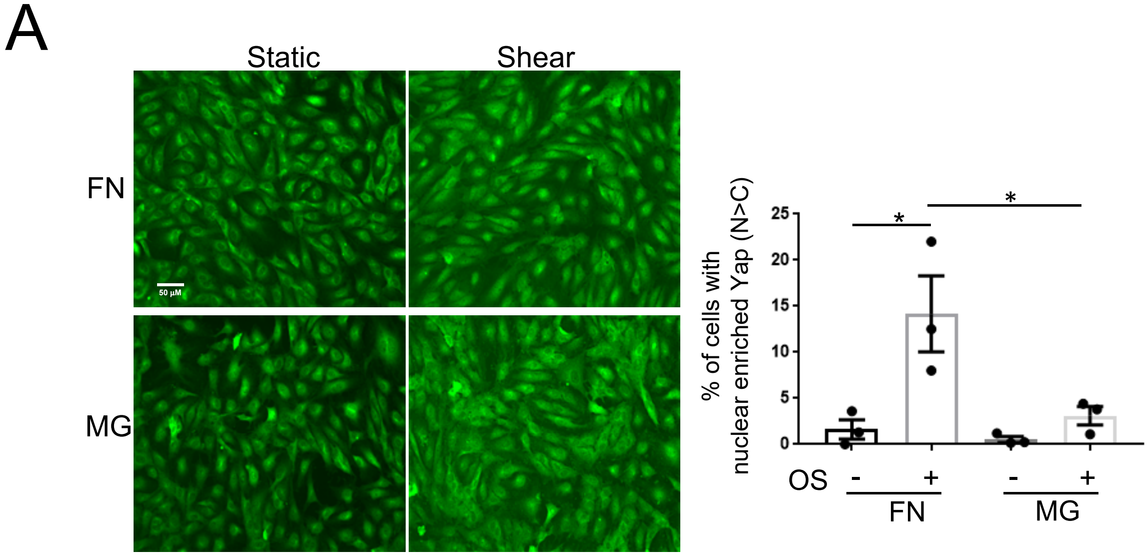


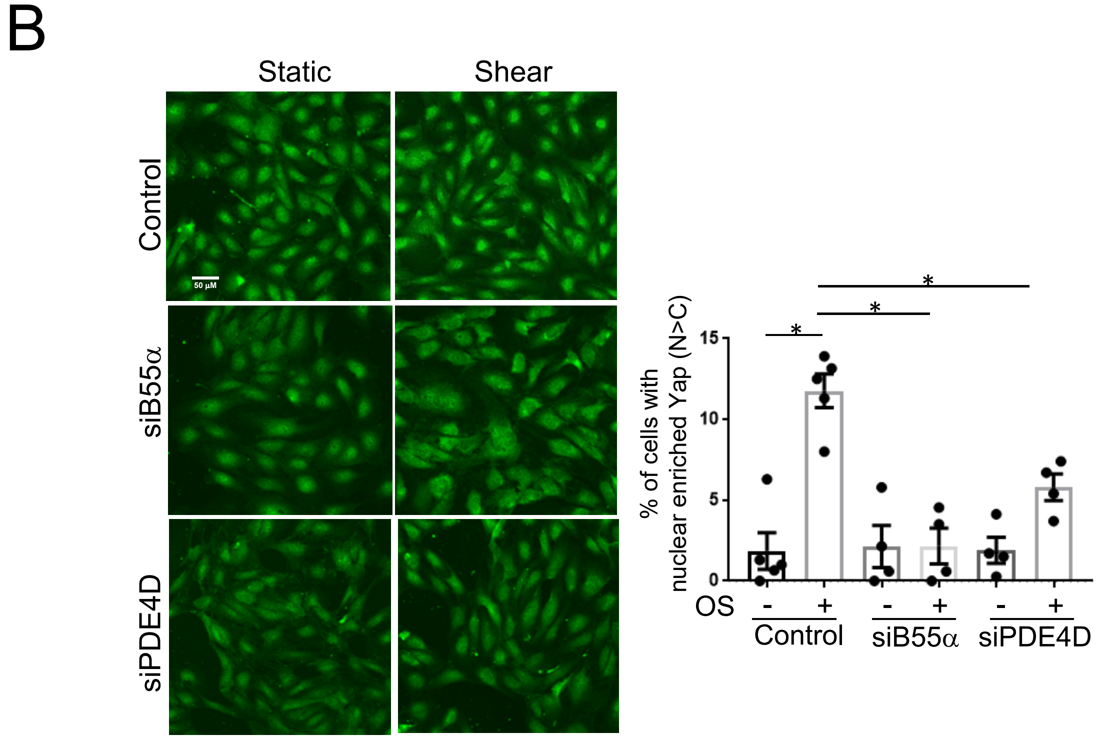


Supplementary Fig 5

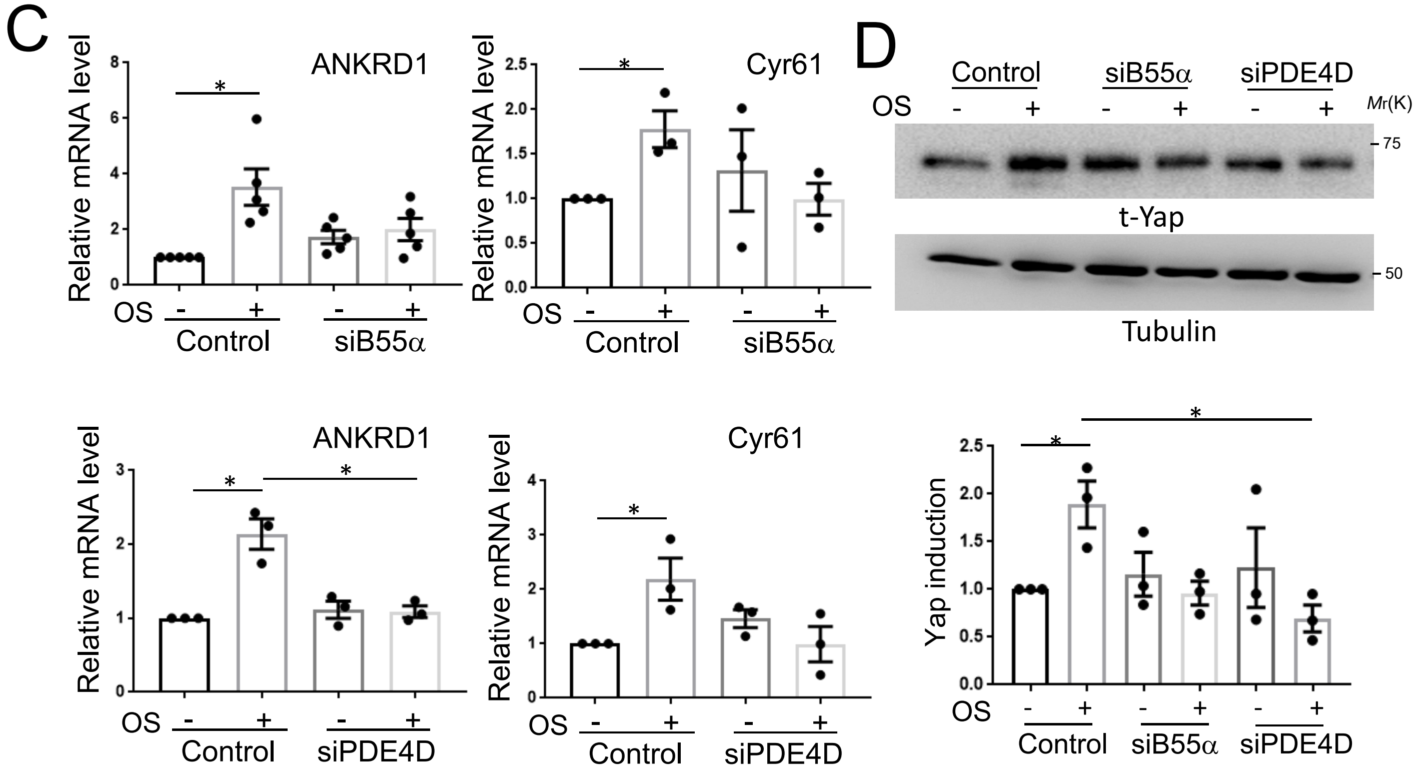


5. Role of PDE4D and B55α on FN-induced Yap activation

**A**. HUVECs were replated on FN or MG for 6 hr and exposed to oscillatory shear for 2 hr, fixed and stained for Yap. Cells with nuclear enriched Yap (N>C) were counted from 10 random fields for each condition (n=3 independent experiments). **B**, HUVECs were transfected with B55α siRNA or PDE4D siRNA and replated on FN, were exposed to oscillatory shear for 2 hr and Yap localization assayed as (A) (n=4-5 independent experiments). **C.** BAECs transfected with B55α siRNA or PDE4D siRNA were subject to oscillatory shear for 18 hr. mRNA was isolated and transcript level of Cyr61 and ANKRD1 was measured using qPCR (n=3). **D**. HUVECs transfected with B55α siRNA or PDE4D siRNA were replated on FN and subjected to oscillatory shear for 18 hr. The cells were lysed and probed for Yap expression (n=3). *$p<0.05$ by two-tailed t-test (C) or one way ANOVA (A, B, D).

Supplementary Fig 5

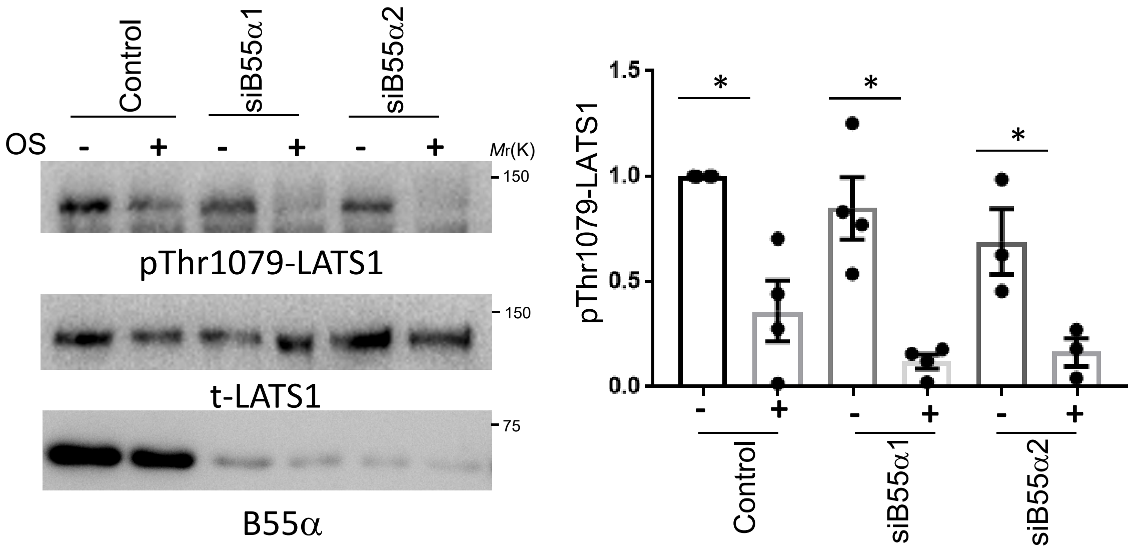


6. B55α regulation of the Hippo pathway

BAECs transfected with B55α siRNAs were plated on FN and subjected to oscillatory shear for 2 hr. Flow-dependent LATS1 phosphorylation was measured using $pT^{1079}$-LATS1 antibody (n=4). *$p<0.05$ by two-tailed t-test.

Supplementary Fig. 6
